# GABAergic neuron-to-glioma synapses in diffuse midline gliomas

**DOI:** 10.1101/2022.11.08.515720

**Authors:** Tara Barron, Belgin Yalçın, Aaron Mochizuki, Evan Cantor, Kiarash Shamardani, Dana Tlais, Andrea Franson, Samantha Lyons, Vilina Mehta, Samin Maleki Jahan, Kathryn R. Taylor, Michael B. Keough, Haojun Xu, Minhui Su, Michael A. Quezada, Pamelyn J Woo, Paul G. Fisher, Cynthia J. Campen, Sonia Partap, Carl Koschmann, Michelle Monje

**Author notes:** Please send correspondence to: Michelle Monje MD PhD.

## Abstract

Pediatric high-grade gliomas are the leading cause of brain cancer-related death in children. High-grade gliomas include clinically and molecularly distinct subtypes that stratify by anatomical location into diffuse midline gliomas (DMG) such as diffuse intrinsic pontine glioma (DIPG) and hemispheric high-grade gliomas. Neuronal activity drives high-grade glioma progression both through paracrine signaling^1,2^ and direct neuron-to-glioma synapses^3–5^. Glutamatergic, AMPA receptor-dependent synapses between neurons and malignant glioma cells have been demonstrated in both pediatric^3^ and adult high-grade gliomas^4^, but neuron-to-glioma synapses mediated by other neurotransmitters remain largely unexplored. Using whole-cell patch clamp electrophysiology, *in vivo* optogenetics and patient-derived glioma xenograft models, we have now identified functional, tumor-promoting GABAergic neuron-to-glioma synapses mediated by GABA_A_ receptors in DMGs. GABAergic input has a depolarizing effect on DMG cells due to NKCC1 expression and consequently elevated intracellular chloride concentration in DMG tumor cells. As membrane depolarization increases glioma proliferation^3^, we find that the activity of GABAergic interneurons promotes DMG proliferation *in vivo*. Increasing GABA signaling with the benzodiazepine lorazepam – a positive allosteric modulator of GABA_A_ receptors commonly administered to children with DMG for nausea or anxiety - increases GABA_A_ receptor conductance and increases glioma proliferation in orthotopic xenograft models of DMG. Conversely, levetiracetam, an anti-epileptic drug that attenuates GABAergic neuron-to-glioma synaptic currents, reduces glioma proliferation in patient-derived DMG xenografts and extends survival of mice bearing DMG xenografts. Concordant with gene expression patterns of GABA_A_ receptor subunit genes across subtypes of glioma, depolarizing GABAergic currents were not found in hemispheric high-grade gliomas. Accordingly, neither lorazepam nor levetiracetam influenced the growth rate of hemispheric high-grade glioma patient-derived xenograft models. Retrospective real-world clinical data are consistent with these conclusions and should be replicated in future prospective clinical studies. Taken together, these findings uncover GABAergic synaptic communication between GABAergic interneurons and diffuse midline glioma cells, underscoring a tumor subtype-specific mechanism of brain cancer neurophysiology with important potential implications for commonly used drugs in this disease context.

## Introduction

Diffuse midline glioma (DMG), which occurs most commonly in the brainstem and is also known as diffuse intrinsic pontine glioma (DIPG), is a lethal childhood central nervous system cancer with few therapeutic options and a median survival of only 10-13 months^6,7^. The majority of DMGs exhibit a mutation in genes encoding histone H3 (H3K27M), and occur in the brainstem, thalamus and spinal cord^8-10^. Multiple lines of evidence support the concept that DMG originates from oligodendroglial lineage precursor cells^11–15^. During postnatal development and adulthood, oligodendroglial precursor cells (OPCs) communicate with neurons through both paracrine factor signaling^16–18^ and through glutamatergic and GABAergic neuron-to-OPC synapses^19–23^; oligodendroglial precursor cell proliferation is robustly regulated by neuronal activity^24^. Similar to these effects on their normal cellular counterparts, glutamatergic neuronal activity drives the proliferation and growth of DMG and other high-grade^1,4,25,26^ and low-grade^2^ gliomas. The mechanisms by which neuronal activity promotes glioma progression include activity-regulated paracrine factor secretion^1,2,25,26^ as well as electrochemical communication through AMPA (α-amino-3-hydroxy-5-methyl-4-isoxazole propionic acid) receptor (AMPAR)-mediated neuron-to-glioma synapses^3,4^ and activity-dependent, potassium-evoked glioma currents that are evident in both pediatric and adult forms of high-grade gliomas^3,4^. Depolarizing current alone is sufficient to drive malignant glioma growth in orthotopic xenograft models ^3^, underscoring the need for a comprehensive understanding of electrochemical mechanisms that enable glioma membrane depolarization in each molecularly distinct form of glioma. Here, we explore whether GABAergic synapses exist between GABAergic interneurons and DMG cells and test the hypothesis that putative GABAergic synaptic signaling is depolarizing and promotes tumor progression in H3K27M-altered DMG.

## Results

### GABA_A_ receptor and postsynaptic gene expression in DMG

To determine whether genes involved in GABAergic synaptic transmission are expressed in high-grade gliomas, we analyzed single-cell RNAseq datasets from primary patient tumor samples of H3K27M+ DMG cells, IDH wild-type (WT) hemispheric high-grade glioma cells, IDH mutant (mut) hemispheric high-grade glioma cells, and tumor-associated non-malignant oligodendrocytes (OLs). H3K27M+ DMG cells broadly expressed GABA_A_ receptor subunit genes, including α and β subunits, as well as ARHGEF9, GPHN, and NLGN2, which are associated with GABAergic post-synaptic regions (Figure 1a). These genes were expressed to a much greater extent in H3K27M+ DMG cells than in IDH WT high-grade gliomas (Figure 1a).

**Figure 1.**
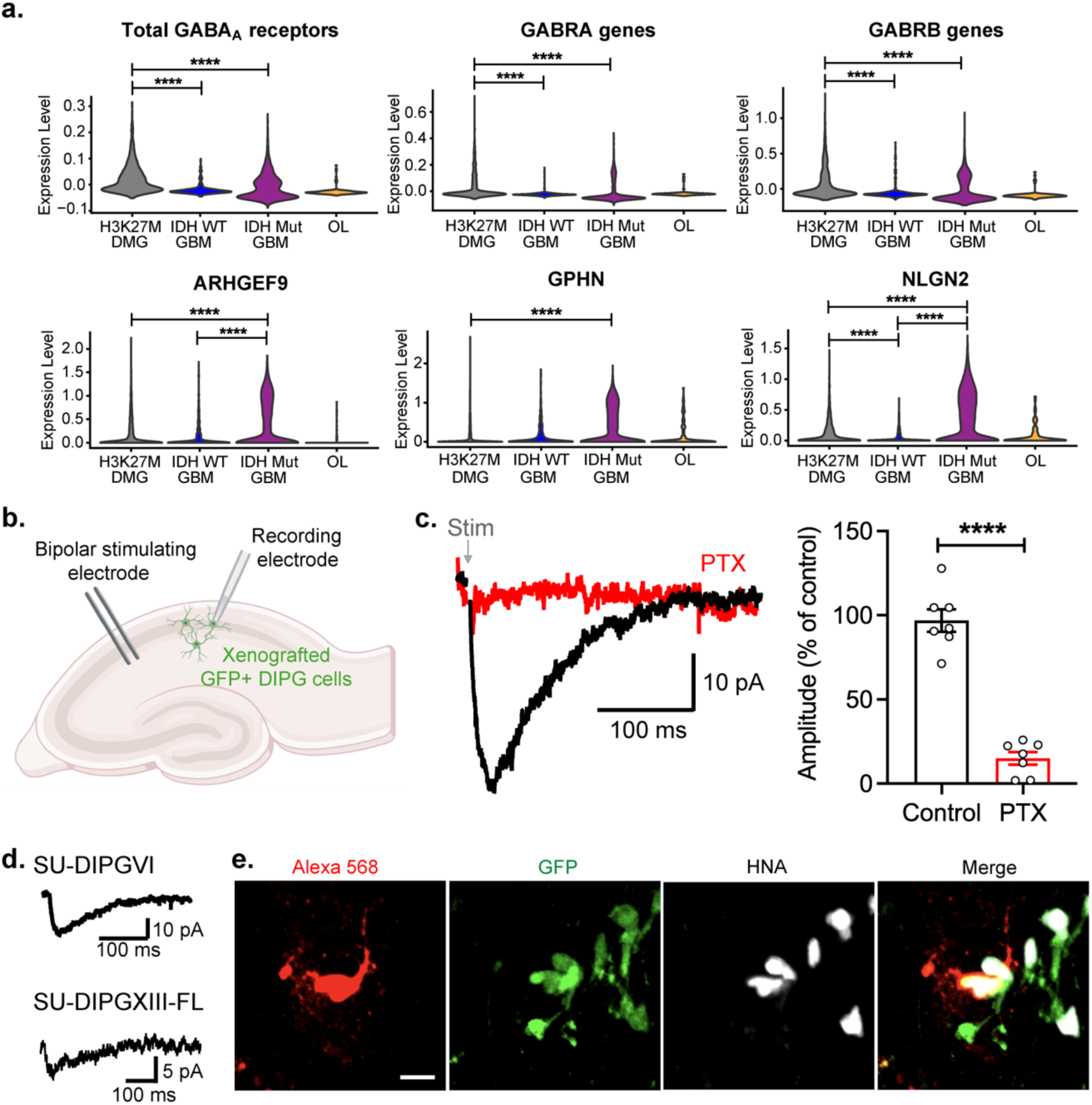
GABAergic neuron-glioma synapses. **a.** Single cell RNAseq analysis of primary human biopsies of H3K27M diffuse midline glioma (grey; n = 2,259 cells, 6 study participants), IDH wild-type (WT) high-grade glioma (blue; n = 599 cells, 3 participants), IDH mutant (mut) high-grade glioma (purple; n = 5,096 cells, 10 participants) malignant cells, and tumor-associated, non-malignant oligodendrocytes (OL, yellow; n = 232 cells), demonstrating expression of total GABA_A_ receptor subunit genes, α subunit genes, β subunit genes, and postsynaptic genes specific to GABAergic synapses. Statistical analyses performed on single cells are represented with stars only when also significant when analyzed on a per patient basis as well as a per cell basis. Comparisons to OL (control cell type) are not shown. **b.** Patient-derived DMG cells expressing GFP were xenografted into the CA1 region of the hippocampus of NSG mice. Response to local CA1 stimulation via a bipolar stimulator was recorded in xenografted cells using whole-cell patch clamp electrophysiology. **c.** Representative trace of picrotoxin (PTX)-sensitive GABAergic postsynaptic current (PSC) in a DMG cell (left). Quantification of current amplitude after 50 µM PTX as a % of control (right; n = 7 cells from 5 mice). Recording performed in the presence of NBQX to block AMPAR currents. **d.** Representative traces of GABAergic PSCs in two xenografted DMG cell lines. **e.** Confocal image of a xenografted DMG cell dye-filled (Alexa 568; red) during recording and co-labelled with GFP (green) and HNA (white) post-recording. Scale bar, 10 µm. All data are mean ± s.e.m. ****P < 0.0001, paired Student’s t-test.

### Functional GABAergic neuron-glioma synapses

To determine whether putative functional GABAergic neuron-glioma synapses exist, we performed electrophysiological recordings from xenografted H3K27M+ DMG cells in response to stimulation of local GABAergic interneurons. Green fluorescent protein (GFP)-expressing glioma cells were xenografted into the CA1 region of the hippocampus, a well-defined circuit in which neuron-to-glioma synapses have been previously reported^3^ and allowed to engraft and grow for at least 8 weeks. Acute hippocampal sections were prepared from these mice, and DMG cell responses to electrical stimulation of local neurons with a bipolar stimulator were recorded using whole-cell patch clamp electrophysiology (Figure 1b). Using a high Cl^-^ internal solution and in the presence of AMPAR antagonist NBQX (2,3-dihydroxy-6-nitro-7-sulfamoyl-benzo[f]quinoxaline) to inhibit AMPAR-mediated currents, local stimulation led to an inward current in DMG cells recorded in voltage clamp (Figure 1c). This current was blocked with perfusion of picrotoxin (PTX), a GABA_A_ receptor inhibitor, indicating that these synaptic currents are mediated by GABA. GABAergic neuron-to-glioma synapses were observed in two distinct patient-derived xenograft models (Figure 1d). Cells were filled with Alexa Fluor 568 dye during electrophysiological recording and then slices were post-fixed, immunostained for tumor cell markers [GFP and human nuclear antigen (HNA)] and imaged on a confocal microscope to confirm that the cells recorded from were glioma cells (Figure 1e).

### GABA depolarizes DMG cells due to NKCC1-mediated high intracellular Cl^-^

GABA_A_ receptor activation can either depolarize or hyperpolarize a cell, depending on the intracellular Cl^-^ concentration. In mature neurons, Cl^-^ concentration is low, leading to an influx of Cl^-^ through GABA_A_ receptors and thus, hyperpolarization ^27^. OPCs exhibit a high intracellular Cl^-^ concentration, leading to an efflux of Cl^-^ through GABA_A_ receptors, and thus GABAergic neuron-to-OPC synapses cause depolarization^20^. Perforated patch recordings of xenografted H3K27M+ DMG cells using gramicidin-A revealed that local application of GABA induced an inward current in voltage clamp and corresponding depolarization in current clamp (Figure 2a-b). The effect of local GABA application on H3/IDH WT pediatric hemispheric glioblastoma (pGBM) was negligible in comparison (Figure 2a-b). PTX inhibited the current and depolarization in response to GABA application, indicating that GABA_A_ receptors are responsible for this depolarizing effect. To determine the reversal potential of GABA_A_ currents (*E*_GABA_) in H3K27M+ DMG and H3/IDH WT pGBM xenografts, we recorded response to local GABA application at varying holding potentials (Figure 2c). The currents in response to GABA were plotted against the holding potentials, and *E*_GABA_ was found to be -19.61 ± 8.29 mV and -14.14 ± 9.04 mV for H3K27M+ DMG cells from two different patient-derived models (SU-DIPGVI and SU-DIPGXIII-FL, respectively) and -47.44 ± 7.79 mV for H3/IDH WT pGBM (Figure 2d; Extended Data Figure 1). Using these reversal potentials to calculate the intracellular Cl^-^ concentration of each glioma type, we find the intracellular Cl^-^ concentration of H3K27M+ DMG cells to be 62.65 mM in SU-DIPGVI cells and 76.88 mM in SU-DIPGXIII-FL cells, and that of H3/IDH WT pGBM to be 22.12 mM. During whole-cell recordings with high Cl^-^ internal solution, the reversal potential was -0.41 ± 12.98 mV in H3K27M+ DMG cells, illustrating the critical role of chloride concentration gradients.

**Figure 2.**
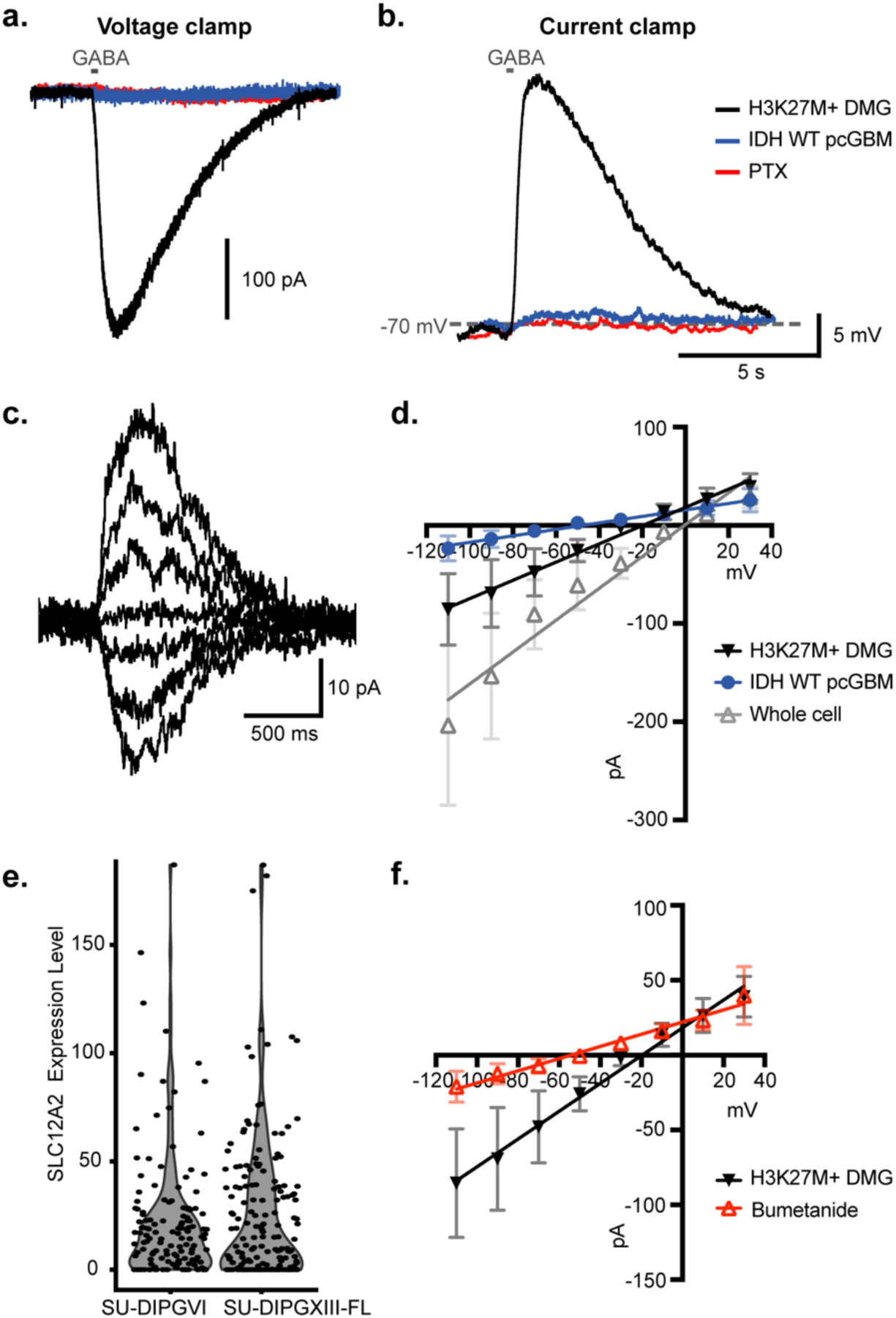
GABA is depolarizing in DMG, but not IDH wild type glioblastoma. **a-b.** Perforated patch of xenografted patient-derived H3K27M+ DMG cells and hemispheric (IDH/H3 WT) pediatric cortical glioblastoma (pcGBM) reveals varying current sizes (in voltage clamp, **a.**) and levels of depolarization (in current clamp, **b.**) in response to local GABA application. **c.** Representative trace of H3K27M+ DMG cell response to GABA at varying membrane potentials. **d.** Current-voltage relationship of GABA current in DMG cells and IDH WT pcGBM cells recorded with perforated patch and whole-cell patch clamp electrophysiology. Reversal potential of GABA was -19.61 mV in H3K27M+ DMG cells (n = 6 cells from 5 mice), -47.44 mV in IDH WT pcGBM cells (n = 6 cells from 5 mice), and -0.4051 mV during whole-cell recording of H3K27M+ DMG cells with a high Cl^-^ internal solution (n = 4 cells from 4 mice). **e.** Single cell RNAseq analysis of SLC12A2 (NKCC1) in patient-derived DMG xenografts. **f.** Current-voltage relationship of H3K27M+ DMG cells in the presence of 100 μM bumetanide, a NKCC1 inhibitor. Reversal potential of GABA in DMG cells is -54.20 mV in the presence of bumetanide (n = 5 cells from 3 mice). All data are mean ± s.e.m.

Cation–chloride cotransporters, such as the Na-K-Cl cotransporter NKCC1, have an important role in setting intracellular Cl^-^ concentration. *SLC12A2*, the gene that encodes for NKCC1, is expressed in H3K27M+ DMG (Figure 2e). To determine the role of NKCC1 in *E*_GABA_ in DMG cells, we used perforated patch to record the response to local GABA application in the presence of bumetanide, an NKCC1 inhibitor. After bath perfusion of bumetanide, *E*_GABA_ was shifted from -19.61 ± 8.29 mV to -54.20 ± 8.19 mV in H3K27M+ DMG cells (Figure 2f), a value similar to that found in H3/IDH WT pGBM cells (Figure 2d) indicating that NKCC1 function is critical for the depolarizing effect of GABA on these cells.

### GABAergic interneurons increase DMG proliferation

Past work has demonstrated that glutamatergic neuronal activity promotes glioma progression^1–5^, and that depolarization of glioma cells plays a central role in these effects of neuronal activity on glioma proliferation^3^. Since GABA has a depolarizing effect on DMG cells as described above, we sought to determine whether GABAergic interneurons drive DMG proliferation through depolarizing GABAergic synaptic input. We first used whole-cell patch clamp electrophysiology to confirm that we could perform optogenetic and pharmacological targeting of GABAergic neuron-to-glioma synapses. We genetically expressed ChRmine, a red-shifted channelrhodopsin^28^, in Dlx-expressing GABAergic interneurons in the CA1 region of the hippocampus and recorded the response of xenografted glioma cells to 5 ms optogenetic stimulation of those neurons (Figure 3a). PTX-sensitive GABAergic post-synaptic currents in DMG cells were observed in response to optogenetic interneuron stimulation (Figure 3b). We also observed the tetrodotoxin (TTX)-sensitive prolonged currents, evoked by activity-dependent extracellular K^+^ increase, that we have previously described^3^ (Figure 3b). Whole cell patch clamp recordings of Dlx-ChRmine-expressing interneurons confirmed that optogenetic stimulation evoked depolarization (Extended Data Figure 2a). Pharmacological targeting of GABAergic neuron-to-glioma synaptic input using a benzodiazepine, lorazepam, which increases conductance of GABA_A_ receptors, increased the amplitude of GABAergic post-synaptic currents in DMG cells (Figure 3c-e).

**Figure 3.**
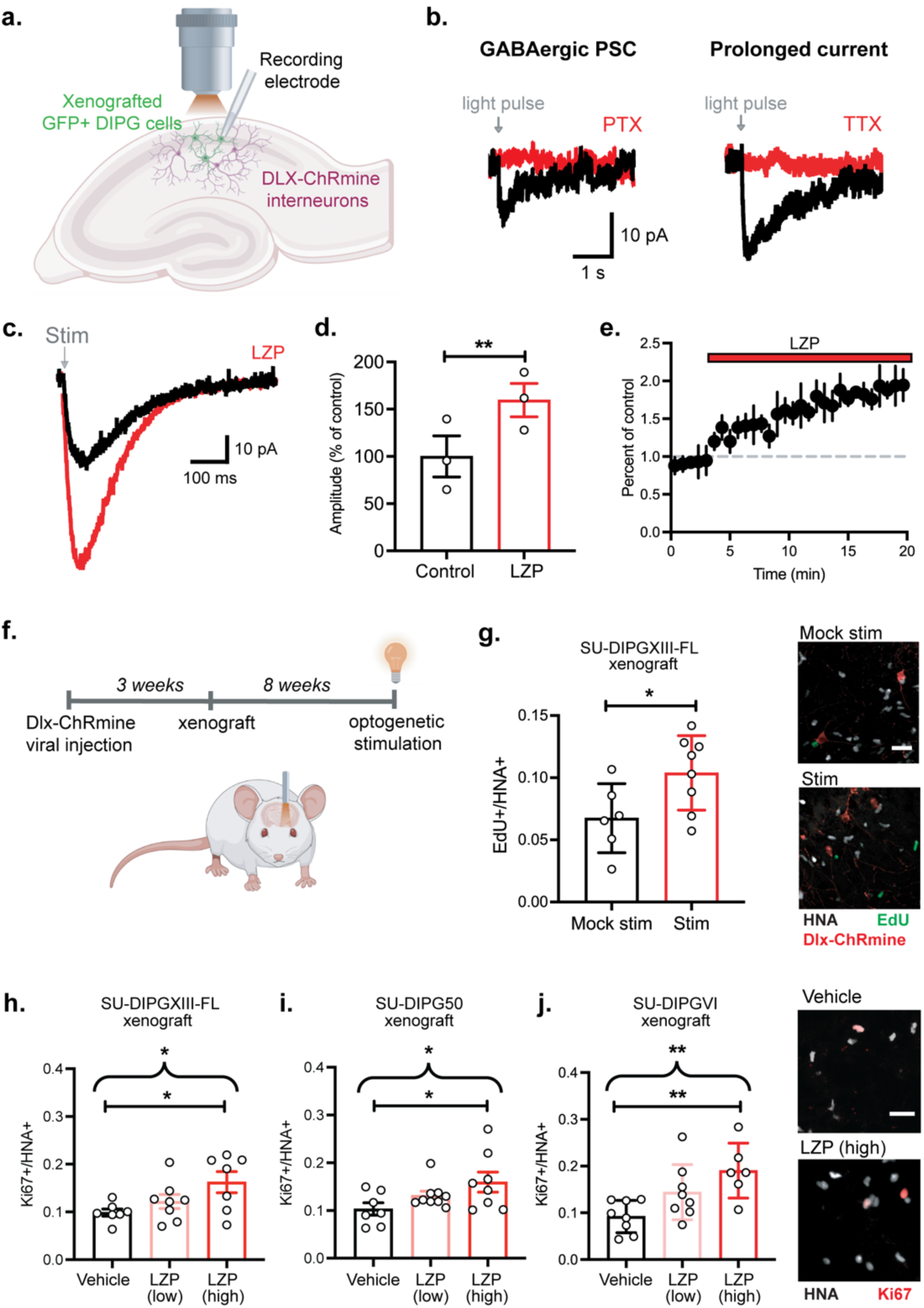
GABAergic interneuron activity drives DMG proliferation. **a.** Patient-derived DMG cells expressing GFP were xenografted into the CA1 region of the hippocampus of NSG mice. Response to optogenetic stimulation of GABAergic interneurons expressing DLX-ChRmine was recorded in xenografted cells using patch clamp electrophysiology. **b.** Two types of responses to optogenetic stimulation of GABAergic neurons were recorded in DMG cells: a PTX-sensitive GABAergic PSC (top) and a prolonged tetrodotoxin (TTX)-sensitive current (bottom). **c.** Representative trace of GABAergic PSCs in DMG in the absence and presence of 10 μM lorazepam (LZP), a benzodiazepine. **d.** Quantification of current amplitude after LZP perfusion as a % of control (n = 3 cells from 3 mice), paired Student’s t-test. **e.** Time course of GABAergic PSC decrease in response to LZP. **f.** Experimental paradigm for *in vivo* optogenetic stimulation of DLX-ChRmine interneurons near xenografted DMG cells in the CA1 region of the hippocampus. **g.** Quantification of proliferation index (EdU+/HNA+ cells) after optogenetic stimulation or mock stimulation (left; mock stim, n = 6 mice; stim, n = 8 mice, two-tailed Student’s t-test). Right, representative confocal images of DLX-ChRmine GABAergic interneurons (red) near xenografted DMG cells expressing EdU (green) and HNA (white). Scale bar, 25 µm. **h-j.** Dose-dependent (low = 2 mg/kg; high = 8 mg/kg) effect of LZP treatment in mice with patient-derived DMG xenografts, SU-DIPGXIII-FL (vehicle, n = 7 mice; low dose, n = 8 mice; high dose, n = 7 mice; **h**), SU-DIPG50 (vehicle, n = 7 mice; low dose, n = 9 mice; high dose, n = 8 mice; **i**), and SU-DIPGVI (vehicle, n = 8 mice; low dose, n = 8 mice; high dose, n = 6 mice; **j**), one-way ANOVA. Straight brackets indicate Dunnett’s multiple comparisons test between two groups; curved brackets indicate post-test for linear contrast among all three groups. Right, representative confocal images of xenografted SU-DIPGVI cells expressing Ki67 (red) and HNA (white). Scale bar, 25 µm. All data are mean ± s.e.m. *P < 0.05, **P < 0.01.

We next sought to test the effect of interneuron activity and GABAergic synaptic input into DMG cells *in vivo*. Dlx-ChRmine was expressed in hippocampal interneurons via AAV viral vector injection to the CA1 region, and *in vivo* optogenetic stimulation of interneuron activity was confirmed by expression of the immediate early gene cFos (Extended Data Figure 2b). Eleven weeks after injection of Dlx-ChRmine vector into the hippocampus, and eight weeks after xenografting patient-derived H3K27M+ DMG cells to the same area, the CA1 region of the hippocampus was optogenetically stimulated (595 nm light, 40 Hz, 30 sec on/90 sec off over 30 minutes) in awake, behaving mice to stimulate GABAergic interneuron activity (Figure 3f). Control mice were identically manipulated, but light was not delivered during mock optogenetic stimulation. The thymidine analogue EdU was administered systemically to mice at the time of optogenetic or mock stimulation to label proliferating cells, and glioma cell proliferation was analyzed 24-hours later. *In vivo* optogenetic stimulation of GABAergic interneurons promoted proliferation of xenografted DMG cells (Figure 3g). Similarly, treatment of xenografted mice with lorazepam, which increases GABA_A_ receptor signaling, exerted a dose-dependent proliferative effect on H3K27M+ DMG in each of three independent patient-derived orthotopic xenograft models (Figure 3h-j). While the effect of lorazepam was most robust at high doses (8 mg/kg), in each xenograft model a significant dose-dependency was evident with ANOVA post-test for linear contrast. The microenvironment of the brain, such as the presence of GABAergic neurons, is required for the proliferative effect of lorazepam, as no effect of lorazepam was observed in H3K27M+ DMG monocultures (Extended Data Figure 3a). As expected, given the lack of GABA-induced currents in H3/IDH WT gliomas, lorazepam did not increase glioma proliferation in mice bearing patient-derived H3/IDH WT pGBM xenografts (Extended Data Figure 4).

**Figure 4.**
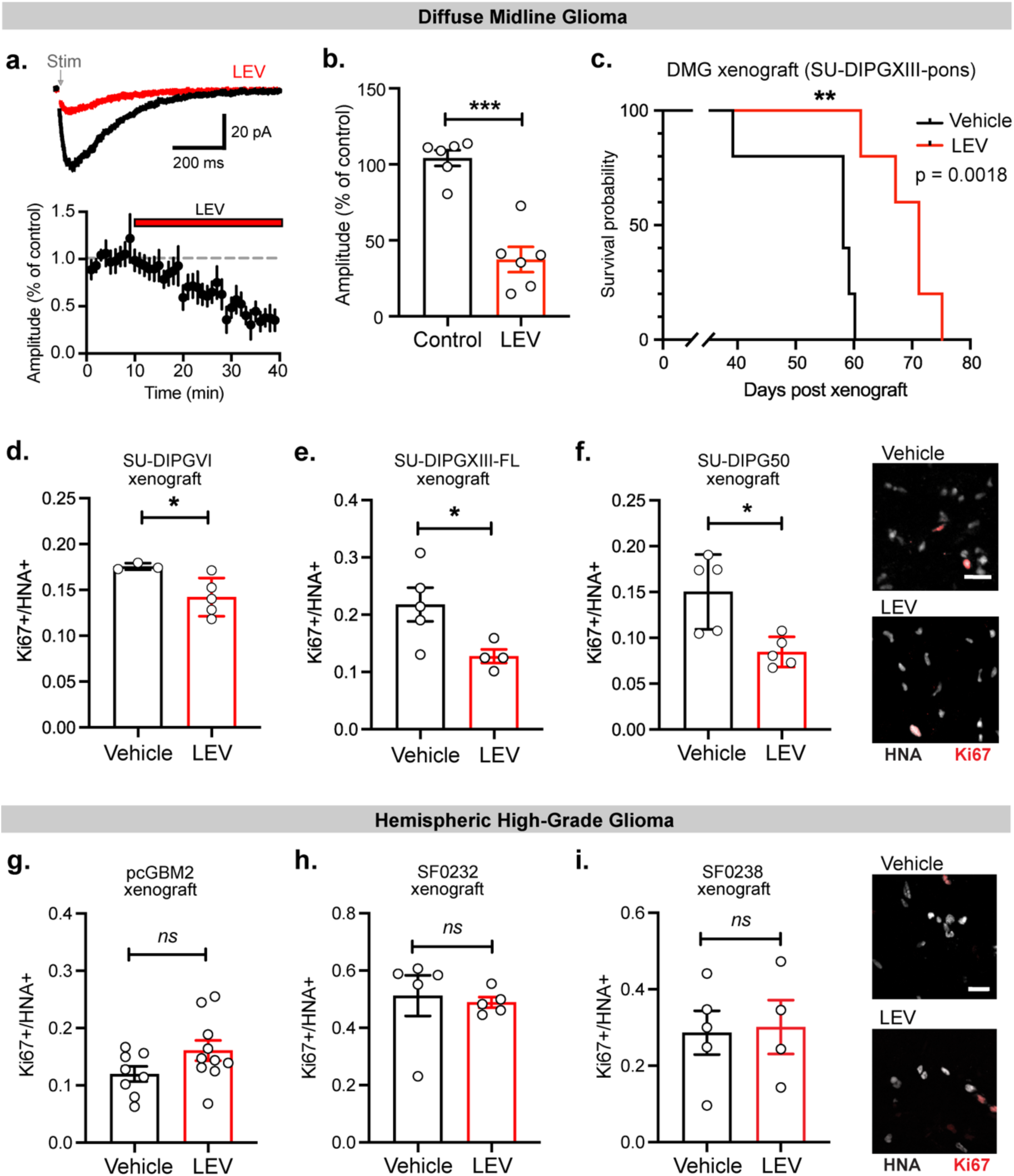
Targeting neuron-to-glioma synapses reduces tumor progression in DMG. **a.** Representative trace of GABAergic PSCs in DMG in the absence and presence of 100 μM levetiracetam (LEV), an anti-epileptic drug. Below, time course of GABAergic PSC decrease in response to LEV. **b.** Quantification of current amplitude after LEV perfusion as a % of control (n = 6 cells from 5 mice), paired Student’s t-test. **c.** Kaplan–Meier survival curves of mice with xenografted SU-DIPG-XIII-P tumors treated with LEV or vehicle (n = 5 mice per group). **d-f.** Effect of LEV treatment in mice with patient-derived DMG xenografts, SU-DIPGVI (vehicle, n = 3; LEV, n = 5; **d.**), SU-DIPGXIII-FL (vehicle, n = 5 mice; LEV, n = 4 mice; **e.**), and SU-DIPG50 (vehicle, n = 5 mice; LEV, n = 5 mice; **f.**), two-tailed Student’s t-test. Right, representative confocal images of xenografted SU-DIPG50 cells expressing Ki67 (red) and HNA (white). Scale bar, 25 µm. **g.-i.** No effect of LEV treatment in mice with patient-derived hemispheric high-grade glioma xenografts, pcGBM2 (vehicle, n = 8; LEV, n = 10; **g.**), SF0232 (vehicle, n = 5 mice; LEV, n = 5 mice; **h.**) and SF0238 (vehicle, n = 5 mice; LEV, n = 4 mice), two-tailed Student’s t-test. Right, representative confocal images of xenografted SF0238 cells expressing Ki67 (red) and HNA (white). Scale bar, 25 µm. All data are mean ± s.e.m. *P < 0.05, **P < 0.01, ***P < 0.001.

### Therapeutic potential of targeting GABAergic neuron-glioma synapses

As neuron-glioma synapses robustly promote glioma cell proliferation and tumor progression, identifying pharmacological treatments that target these synapses has high therapeutic potential. Levetiracetam, a generally well-tolerated anti-epileptic drug with multiple mechanisms of action, reduces GABAergic post-synaptic currents in DMG cells (Figure 4a-b). Strikingly, mice bearing H3K27M+ DMG xenografts treated with levetiracetam exhibited longer survival than vehicle-treated controls (Figure 4c). Levetiracetam treatment decreased glioma proliferation in mice bearing H3K27M+ DMG xenografts compared to vehicle-treated controls, an effect observed in three independent patient-derived orthotopic xenograft models of H3K27M+ DMG (Figure 4d-f). The effect of levetiracetam on glioma proliferation is dependent on the brain microenvironment rather than cell-intrinsic effects, as no effect of levetiracetam was observed in H3K27M+ DMG monocultures (Extended Data Figure 3b).

Retrospective, real-world data from two major US pediatric neuro-oncology centers (Stanford University and University of Michigan) was assessed to query possible effects of levetiracetam on overall survival in pediatric patients with high-grade gliomas. Kaplan-Meier analysis of all pHGG patients (n = 216) suggests a survival advantage of levetiracetam usage (Extended Data Figure 5a). For multivariable survival analysis, we utilized an elastic net-regularized Cox regression for variable selection and found that in all patients with high-grade glioma, a diagnosis of DMG was – as expected - associated with decreased overall survival (coefficient +0.55), and that thalamic DMG tumor location (coefficient -0.20), and levetiracetam (coefficient -0.11) were associated with increased overall survival; the variables of age, sex, ONC201 usage, and panobinostat usage had coefficients of zero. Conventional or targeted chemotherapy other than ONC201 and panobinostat were surprisingly associated with increased overall survival (coefficient - 0.61), which may be explained by the observation that pontine DMG subjects in the historical database often did not receive any conventional or targeted therapy (Extended Data Table 1) due to demonstrated lack of efficacy of conventional chemotherapy in pontine DMG^29^. Hypothesizing that DMGs drove the positive survival association of levetiracetam usage, we next evaluated DMG and hemispheric HGGs separately. These databases include subjects with biopsy-demonstrated H3K27M-mutated or H3WT diffuse midline gliomas as well as subjects prior to availability of molecular testing for whom diagnosis was based only on the typical radiographic appearance of DMGs; both H3K27M-altered and H3WT subgroups of DMGs are therefore included (Extended Data Table 1). The DMG analysis suggests that patients with DMG who had a history of levetiracetam usage (n = 15 children) exhibited a longer median overall survival (OS) compared to those without levetiracetam usage (n = 105 children; Extended Data Figure 5b, Extended Data Table 1). Those DMG patients with a history of levetiracetam usage had a median OS of 20.3 months, compared to those without levetiracetam usage who exhibited a median OS of 9.2 months (P=0.025). Of note, thalamic DMG represented a higher proportion of the group with a history of levetiracetam usage than the group with no history of levetiracetam usage. Comparing subjects with thalamic and pontine DMG who received levetiracetam, we find no difference in OS in this levetiracetam usage group (Extended Data Figure 5c), suggesting that the higher proportion of thalamic DMG in this group does not account for the observed increased median OS compared to the group without levetiracetam usage. The median OS of the control (no levetiracetam usage) group is consistent with the expected median OS for DIPG/DMG (10-11 months for pontine DMG, 13 months for thalamic DMG)^6,7^. Important caveats are that these data are retrospective, the numbers are small, and levetiracetam should be studied in future prospective clinical studies with stratification by molecular subtype and DMG location before drawing conclusions.

In contrast to the anti-proliferative effect of levetiracetam on xenografted H3K27M+ DIPG/DMG, levetiracetam treatment did not significantly affect glioma proliferation in mice bearing patient-derived hemispheric (H3/IDH WT) high-grade glioma xenografts in three independent models of pediatric and adult hemispheric H3/IDH WT glioblastoma (Figure 4g-i). Concordantly, analysis of the retrospective clinical data focused on hemispheric pediatric high-grade gliomas revealed no effect of levetiracetam usage on OS in pediatric patients with non-DMG, hemispheric high-grade gliomas (n=37, median OS 24.6 months vs n=60 children, median OS 17.0 months with and without levetiracetam use, respectively, P=0.74, Extended Data Figure 5d, Extended Data Table 2).

Phenytoin and ethosuximide, antiepileptic drugs that reduce neuronal hyperexcitability but do not directly act on known mechanisms of neuron-to-glioma communication, do not influence DMG proliferation *in vivo* or *in vitro* (Extended Data Figure 6), highlighting the importance of specifically targeting neuron-to-glioma synapses.

## Discussion

Glutamatergic neuronal activity has emerged as a powerful regulator of glioma progression^2–4,25,26,30^. Across multiple clinically and molecularly distinct forms of pediatric and adult gliomas, activity-regulated paracrine factors such as BDNF and shed neuroligin-3 promote glioma growth ^1,2,25,30^. Similarly, AMPAR-mediated glutamatergic synapses drive progression in both H3K27M-altered DMG and hemispheric (H3/IDH WT) glioblastomas^3,4^. Here, we demonstrate that GABAergic interneurons also promote glioma progression through GABAergic synapses that are depolarizing and growth-promoting in the specific disease context of diffuse midline gliomas. In contrast, only minimal currents were found in the hemispheric (IDH/H3 WT) high-grade glioma models used here; it is possible that some subtypes of hemispheric glioma may be found to respond to GABA heterogenously^31^. We found that the commonly used anti-seizure drug levetiracetam attenuates these GABAergic currents in diffuse midline gliomas, through mechanisms that remain to be determined. Levetiracetam has multiple described mechanisms, including binding to SV2A to decrease presynaptic release, but whether this represents the mechanism operant in diminishing DMG GABA currents remains to be tested in future studies. These discoveries highlight the therapeutic potential of re-purposing levetiracetam to decrease GABAergic signaling in diffuse midline gliomas.

While this therapeutic potential is supported by the preclinical evidence and suggested by the retrospective clinical data presented here, it is important to note that prospective clinical trials are required to validate this effect. In this clinical retrospective study, it may be that those subjects who developed seizures and therefore received levetiracetam had tumors that are particularly neurotrophic and thus more susceptible to therapy with levetiracetam. It is also possible that tumors growing in neuroanatomical locations with relatively more GABAergic input or different GABA-dependent circuit dynamics are differentially affected by levetiracetam therapy. An unknown confounder could also be associated with levetiracetam therapy. Future work, studying larger numbers of patients and stratifying subjects based on DMG location, molecular characteristics, and glioma neuroscience correlative markers will be required to draw conclusions about the potential role of levetiracetam for DMG therapy.

The anti-seizure drugs tested were growth-inhibitory only if the drug targeted specific mechanisms of neuron-glioma interaction in that tumor type. Ethosuximide and phenytoin do not target known mechanisms of neuron-glioma interactions and did not affect tumor proliferation in the preclinical models used here. Similarly, neither levetiracetam nor lorazepam influenced the proliferation of the three independent hemispheric (H3/IDH WT) glioblastoma models used here. Past clinical studies of antiepileptic drug effects in adult high-grade glioma have not been guided by knowledge of drugs that specifically target neurophysiological mechanisms operant in that tumor type. Not surprisingly, the results of such anti-seizure medication studies have been mixed. Levetiracetam used concomitantly with chemoradiation has been reported to improve outcomes in hemispheric, H3/IDH WT glioblastoma in some studies ^32^, while large meta-analyses have found no discernable effect on outcome in others ^33,34^. These discordant findings in the literature may reflect the heterogeneity inherent in hemispheric H3/IDH WT high-grade gliomas^34^, and specific subgroups of H3/IDH WT glioblastoma yet-to-be determined could be responsive to levetiracetam. Here, we found no effect of levetiracetam in three independent preclinical models of pediatric and adult hemispheric (H3/IDH WT) high-grade glioma and no effect of levetiracetam in retrospective analyses of pediatric patients with non-DMG, hemispheric high-grade gliomas.

In DMGs, the risk to benefit ratio of benzodiazepines should be carefully considered. Benzodiazepines, which potentiate signaling through GABA_A_ receptors, promote glioma GABAergic currents and tumor proliferation in the H3K27M-altered DMG models used here. Benzodiazepines are commonly used in children with DMG for nausea, anxiety, claustrophobia during MRI scans and other medical procedures, and for other reasons. While benzodiazepines are important medications for palliative care, use should be carefully considered in DMG outside of the context of end-of-life care and should be further evaluated in clinical analyses. Conversely, and further underscoring differences between DMG and hemispheric high-grade gliomas, preclinical studies indicate that GABA and GABAergic interneurons may instead be growth-inhibitory hemispheric (H3/IDH WT) adult glioblastoma models^35,36^. These findings underscore the therapeutic importance of elucidating the neurophysiology of defined subtypes of brain cancers to identify the patient populations for which a particular neurophysiological drug may be beneficial or detrimental. Understanding the neuroscience of brain tumors will enable the development of effective and safe therapeutic approaches, incorporating neuroscience-informed therapies into combinatorial strategies targeting both cell-intrinsic and microenvironmental mechanisms that drive progression of these devastating cancers.

## Methods

### Human samples and data

For all human tissue and cell studies, informed consent was obtained, and tissue was used in accordance with protocols approved by the Stanford University Institutional Review Board (IRB). IRB approval was also obtained for retrospective analyses of real-world clinical data kept in IRB-approved databases at Stanford University and University Michigan.

### Mice and housing conditions

All *in vivo* experiments were conducted in accordance with protocols approved by the Stanford University Institutional Animal Care and Use Committee (IACUC) and performed in accordance with institutional guidelines. Animals were housed according to standard guidelines with free access to food and water in a 12 h light:12 h dark cycle. For brain tumor xenograft experiments, the IACUC does not set a limit on maximal tumor volume but rather on indications of morbidity. In no experiments were these limits exceeded as mice were euthanized if they exhibited signs of neurological morbidity or if they lost 15% or more of their body weight.

### Orthotopic xenografting

For all xenograft studies, NSG mice (NOD-SCID-IL2R gamma chain-deficient, The Jackson Laboratory) were used. Male and female mice were used equally. A single-cell suspension from cultured SU-DIPG-VI-GFP, SU-DIPG-XIII-FL-GFP, SU-DIPG-50-GFP, SU-pcGBM2-GFP, SF0232, or SF0238 neurospheres were prepared in sterile PBS immediately before the xenograft procedure. Animals at postnatal day (P) 28–30 were anaesthetized with 1–4% isoflurane and placed in a stereotactic apparatus. The cranium was exposed via midline incision under aseptic conditions. Approximately 300,000 cells in 3 µl sterile PBS were stereotactically implanted through a 26-gauge burr hole, using a digital pump at infusion rate of 0.4 µl min^−1^ and 26-gauge Hamilton syringe. For all electrophysiology and optogenetics experiments, cells were implanted into the CA1 region of the hippocampus (1.5 mm lateral to midline, -1.8 mm posterior to bregma, −1.4 mm deep to cranial surface). SU-DIPG-XIII-FL-GFP for lorazepam and levetiracetam treatments were xenografted into the premotor cortex (0.5 mm lateral to midline, 1.0 mm anterior to bregma, −1.75 mm deep to cranial surface). SU-DIPG-XIII-P for survival study and SU-DIPG-VI-GFP and SU-DIPG-50-GFP for lorazepam, levetiracetam, ethosuximide, and phenytoin treatments were xenografted into the pons (1.0 mm lateral to midline, −0.8 mm posterior to lambda, −5.0 mm deep to cranial surface). At the completion of infusion, the syringe needle was allowed to remain in place for a minimum of 2 min, then manually withdrawn at a rate of 0.875 mm min^−1^ to minimize backflow of the injected cell suspension.

### Patient-derived cell culture

All high-grade glioma cultures were generated as previously described ^11^. In brief, tissue was obtained from high-grade glioma (WHO (World Health Organization) grade III or IV) tumors at the time of biopsy or from early post-mortem donations. Tissue was dissociated both mechanically and enzymatically and grown in a defined, serum-free medium designated ‘tumor stem media’ (TSM), consisting of neurobasal(-A) (Invitrogen), B27(-A) (Invitrogen), human bFGF (20 ng ml^−1^; Shenandoah), human EGF (20 ng ml^−1^; Shenandoah), human PDGF-AA (10 ng ml^−1^) and PDGF-BB (10 ng ml−1; Shenandoah) and heparin (2 ng ml^−1^; Stem Cell Technologies). For all patient-derived cultures, mycoplasma testing was routinely performed, and short tandem repeat DNA fingerprinting was performed every three months to verify authenticity. The short tandem repeat fingerprints and clinical characteristics for the patient-derived cultures and xenograft models used have been previously reported ^37^.

### Single-cell sequencing analysis

We combined publicly available single-cell datasets processed and annotated previously^13,38^, all sequenced using smart-seq2 protocol. Following the quality-control measures taken in these studies, we filtered the data to keep cells with more than 400 detected genes, and genes that were expressed in more than 3 cells. We assessed the single-cell transcriptome from 6,341 adult IDH-mutant glioma cells derived from biopsies from 10 study participants, 599 adult wild-type IDH glioma cells derived from biopsies from 3 study participants, and 2,458 pediatric H3K27M DMG cells derived from biopsies from 6 study participants, as well as the single-cell transcriptome of patient-derived SU-DIPGVI and SU-DIPGXIII-FL cells. Malignant cells were inferred by expression programs and detection of tumor-specific genetic alterations. For each sample, we performed first cell-level normalization, and then centered the gene expression around 0 to allow principal component analysis (PCA) computation. Following the PCA reduction, we clustered the cells using shared nearest neighbor clustering. To examine the various GABA_A_ receptor signatures of each of the cells in each cluster, we used the function AddModuleScore by Seurat package, which calculates the average expression levels of the gene set subtracted by the aggregated expression of 100 randomly chosen control gene sets, where the control gene sets are chosen from matching 25 expression bins corresponding to the tested gene set expression. The gene sets used are as followed: GABA_A_ receptor α: GABRA1, GABRA2, GABRA3, GABRA4, GABRA5, GABRA6; GABA_A_ receptor β: GABRB1, GABRB2, GABRB3; total GABA_A_ receptor: GABRA1, GABRA2, GABRA3, GABRA4, GABRA5, GABRA6, GABRB1, GABRB2, GABRB3, GABRG1, GABRG2, GABRG3, GABRD, GABRE, GABRP, GABRQ, GABRR1, GABRR2, GABRR3.

### Slice preparation for electrophysiology

Coronal slices (300 µm thick) containing the hippocampal region were prepared from mice (at least 8 weeks after xenografting) in accordance with a protocol approved by Stanford University IACUC. After rapid decapitation, the brain was removed from the skull and immersed in ice-cold slicing artificial cerebrospinal fluid (ACSF) containing (in mM): 125 NaCl, 2.5 KCl, 25 glucose, 25 NaHCO_3_ and 1.25 NaH_2_PO_4_, 3 MgCl_2_ and 0.1 CaCl_2_. After cutting, slices were incubated for 30 min in warm (30 °C) oxygenated (95% O2, 5% CO2) recovery ACSF containing (in mM): 100 NaCl, 2.5 KCl, 25 glucose, 25 NaHCO_3_, 1.25 NaH_2_PO4, 30 sucrose, 2 MgCl_2_ and 1 CaCl_2_ before being allowed to equilibrate at room temperature for an additional 30 min.

### Electrophysiology

Slices were transferred to a recording chamber and perfused with oxygenated, warmed (28–30 °C) recording ACSF containing (in mM): 125 NaCl, 2.5 KCl, 25 glucose, 25 NaHCO_3_, 1.25 NaH_2_PO_4_, 1 MgCl_2_ and 2 CaCl_2_. NBQX (10 µM) was perfused with the recording ACSF to prevent AMPA receptor-mediated currents in synaptic response experiments. TTX (0.5 µM) was perfused with the recording ACSF to prevent neuronal action potential firing in GABA puff experiments. Slices were visualized using a microscope equipped with DIC optics (Olympus BX51WI). Recording patch pipettes (3-5 MΩ) were filled with CsCl-based pipette solution containing (in mM): 150 CsCl, 5 EGTA, 1 MgCl_2_, 10 HEPES, 2 ATP, 0.3 GTP, pH = 7.3. Pipette solution additionally contained Alexa 568 (50 μM) to visualize the cell through dye-filling during whole-cell recordings. Gramicidin A (60 μg/mL) was added to the pipette solution for perforated patch recordings. Glioma cells were voltage-clamped at −70 mV. Synaptic responses were evoked with a bipolar electrode connected to an Iso-flex stimulus isolator (A.M.P.I.) placed near the xenografted cells. GABA (1 mM) in recording ACSF was applied via a puff pipette, which was placed approximately 100 μm away from the patched cell and controlled by a Picospritzer II (Parker Hannifin Corp.). Optogenetic currents were evoked with a 598 nm LED using a pE-4000 illumination system (CoolLED). Signals were acquired with a MultiClamp 700B amplifier (Molecular Devices) and digitized at 10 kHz with an InstruTECH LIH 8+8 data acquisition device (HEKA). Data were recorded and analyzed using AxoGraph X (AxoGraph Scientific) and IGOR Pro 8 (Wavemetrics). For representative traces, stimulus artifacts preceding the synaptic currents have been removed for clarity. Intracellular chloride concentration was calculated using the Nernst equation.

### Inhibitors

Drugs and toxins used for electrophysiology were picrotoxin (50 µM; Tocris), TTX (0.5 µM; Tocris), NBQX (10 µM; Tocris), bumetanide (100 μM), lorazepam (10 μM; Hospira), and levetiracetam (100 μM; Selleck Chemicals). When used for *in vitro* slice application, drugs were made up as a stock in distilled water or dimethylsulfoxide (DMSO) and dissolved to their final concentrations in ACSF before exposure to slices.

### Viral injection and fibre optic placement

Animals were anesthetized with 1-4% isoflurane and placed in a stereotaxic apparatus. For optogenetic stimulation experiments, 1 µl of AAV8-Dlx5/6-ChRmine::oScarlet (virus titer= 1.19×1012) (a gift from Dr. Karl Deisseroth from Stanford University; Chen et al., 2020, Nature Biotech) was unilaterally injected using Hamilton Neurosyringe and Stoelting stereotaxic injector over 5 minutes. The viral vector was injected into hippocampus CA1 in the right hemisphere at coordinates: 1.5 mm lateral to midline, -1.8 mm posterior to bregma, -1.3 mm deep to cranial surface. 2 weeks following the viral injection, SU-DIPG-XIII-FL cells were xenografted as described above. After 7 weeks of tumor engraftment, an optic ferrule was placed above the CA1 of the hippocampus of the right hemisphere, at 1.5 mm lateral to midline, -1.8 mm posterior to bregma, -1.25 mm deep to cranial surface.

### Optogenetic stimulation

Optogenetic stimulations were performed at least 10 weeks after the viral vector delivery, 8 weeks after xenografts, and 1 week after optic ferrule implantation. Freely moving animals were connected to a 595 nm fiber-coupled LED laser system with a monofiber patch cord. Optogenetic stimulation was performed with cycles of 595 nm light pulses at 40 Hz frequency, 10 ms width, and a light power output of 10-15mW from the tip of the optic fiber, which lasted for 30 seconds, followed by 90 seconds recovery over a 30-minute period. Animals were injected intraperitoneally with 40 mg/kg EdU (5-ethynyl-2’-deoxyuridine; Invitrogen, E10187) before the session, and were perfused 24 hours after the optogenetic stimulations.

### Bioluminescence imaging

For in vivo monitoring of tumor growth, bioluminescence imaging was performed using an IVIS imaging system (Xenogen). Mice orthotopically xenografted with luciferase-expressing glioma cells were placed under isofluorane anesthesia and injected with luciferin substrate. Animals were imaged at baseline and randomized based on tumor size by a blinded investigator so that experimental groups contained an equivalent range of tumor sizes. Over the course of each study (described below), all total flux values were then normalized to baseline values to determine fold change of tumor growth.

### Mouse drug treatment studies

For all drug studies, NSG mice were xenografted as above with SU-DIPG-VI-GFP, SU-DIPG-XIII-FL-GFP, SU-DIPG-50-GFP, SU-pcGBM2-GFP, SF0232, or SF0238 cells and randomized to treatment group by a blinded investigator. Four to six weeks post-xenograft, mice were treated with systemic administration of lorazepam (8 mg kg^−1^ or 2 mg kg^−1^; Hospira), levetiracetam (20 mg kg^−1^; Selleck Chemicals), or phenytoin (50 mg kg^−1^; Selleck Chemicals) via intraperitoneal injection, or ethosuximide (300 mg kg^−1^; Selleck Chemicals) via oral gavage for four weeks (5 days per week). For all studies, controls were treated with an identical volume of the relevant vehicle. Bioluminescence imaging was performed before treatment and every 7 days thereafter using an IVIS imaging system (Xenogen) under isoflurane anesthesia. Tumor burden was assessed as fold change in total flux from the beginning to end of treatment.

### Xenograft survival studies

For survival studies, morbidity criteria used were either reduction of weight by 15% initial weight, or severe neurological motor deficits consistent with brainstem dysfunction (that is, hemiplegia or an incessant stereotyped circling behavior seen with ventral midbrain dysfunction). Kaplan–Meier survival analysis using log rank testing was performed to determine statistical significance.

### Perfusion and immunohistochemistry

Animals were anaesthetized with intraperitoneal avertin (tribromoethanol), then transcardially perfused with 20 ml of PBS. Brains were fixed in 4% PFA overnight at 4 °C, then transferred to 30% sucrose for cryoprotection. Brains were then embedded in Tissue-Tek O.C.T. (Sakura) and sectioned in the coronal plane at 40 µm using a sliding microtome (Microm HM450; Thermo Scientific).

For immunohistochemistry, coronal sections were incubated in blocking solution (3% normal donkey serum, 0.3% Triton X-100 in TBS) at room temperature for 2 hours. Chicken anti-GFP (1:500, Abcam), mouse anti-human nuclei clone 235-1(1:100; Millipore), or rabbit anti-Ki67 (1:500; Abcam) were diluted in antibody diluent solution (1% normal donkey serum in 0.3% Triton X-100 in TBS) and incubated overnight at 4 °C. Sections were then rinsed three times in TBS and incubated in secondary antibody solution containing Alexa 488 donkey anti-chicken IgG, Alexa 594 donkey anti-rabbit IgG, or Alexa 647 donkey anti-mouse IgG used at 1:500 (Jackson Immuno Research) in antibody diluent at 4 °C overnight. Sections were rinsed three times in TBS and mounted with ProLong Gold Mounting medium (Life Technologies).

### Confocal imaging and quantification of cell proliferation

Cell quantification within xenografts was performed by a blinded investigator using live counting on a 40× oil immersion objective or 20× air objective of a Zeiss LSM700 or Zeiss LSM800 scanning confocal microscope and Zen imaging software (Carl Zeiss). For Ki67 analysis, 3 fields for quantification were selected from each of 3 consecutive sections in a 1-in-6 series of 40-μm coronal sections with respect to overall tumor burden. Within each field, all HNA-positive and GFP-positive tumor cells were quantified to determine tumor burden within the areas quantified. HNA-positive were then assessed for co-labelling with Ki67. To calculate the proliferation index (the percentage of proliferating tumor cells for each mouse), the total number of HNA-positive cells co-labelled with Ki67 across all areas quantified was divided by the total number of cells counted across all areas quantified (Ki67+/HNA+).

### EdU Incorporation Assay

Diffuse intrinsic pontine glioma (DIPG) tumor neurosphere cultures SU-DIPGVI, SU-DIPGXIII, and SU-DIPG50 were generated as previously described^3,7^ from early post-mortem tissue donations and grown as tumor neurospheres in defined, serum-free ‘tumor stem media’ (TSM) media, consisting of 1:1 mixture of neurobasal(-A) (Invitrogen) and D-MEM/F-12 (Invitrogen), HEPES buffer (Invitrogen), MEM sodium pyruvate (Invitrogen), MEM non-essential amino acids (Invitrogen), GlutaMAX-1 supplement (Invitrogen), B27(-A) (Invitrogen), human bFGF (20 ng/ml; Shenandoah), human EGF (20 ng/ml; Shenandoah), human PDGF-AA (10 ng/ml) and PDGF-BB (10 ng/ml; Shenandoah) and heparin (2 ng/ml; Stem Cell Technologies).

100,000 glioma cells were plated onto circular glass coverslips (Electron Microscopy Services) pre-treated for 1 h at 37 °C with poly-L-lysine (Sigma) and then 1 h at 37 °C with 10 µg/ml natural mouse laminin (Thermo Fisher). Dimethyl sulfoxide (Sigma-Aldrich) or drugs at the concentrations indicated (dissolved in dimethyl sulfoxide) were added to the coverslips. 10 μM EdU was added to each coverslip. Cells were fixed after 24 hr using 4% paraformaldehyde in PBS and stained using the Click-iT EdU kit and protocol (Invitrogen). Proliferation index was then determined by quantifying the fraction of EdU labeled cells/DAPI labeled cells using confocal microscopy.

### Retrospective, real-world patient data

Retrospective data on patients with high-grade glial tumors were collected from patient databases at Stanford University (1990-2020) and the University of Michigan (2012-2021) through protocols approved by the respective institutional review boards. Database source data for pediatric high-grade glioma patients were reviewed for this study to ensure veracity and completeness. Overall survival was calculated using the Kaplan-Meier estimator; the log-rank test was utilized to compare survival distributions. Patients were censored at time of last contact for the Kaplan-Meier analysis. Given the number of potential parameters with high correlation, an elastic net-regularized regression was utilized for covariate selection in a multivariable survival model. Clinical data including age; sex; tumor location; diagnosis of DMG; and administration of ONC201, panobinostat, conventional chemotherapy, and levetiracetam were considered potential covariates. Twenty-fold cross-validation was used to obtain the value of λ that gave the minimum mean cross-validated error; corresponding coefficients for each variable were subsequently determined. All data were compiled and analyzed in R version 4.0 or higher.

### Statistical analyses

Statistical tests were conducted using Prism (GraphPad) software unless otherwise indicated. Gaussian distribution was confirmed by the Shapiro–Wilk normality test. For parametric data, unpaired two-tailed Student’s t-tests or one-way ANOVA with Tukey’s post hoc tests to examine pairwise differences were used as indicated. Paired two-tailed Student’s t-tests were used in the case of same cell experiments (as in electrophysiological recordings). For non-parametric data, a two-sided unpaired Mann– Whitney test was used as indicated, or a one-tailed Wilcoxon matched-pairs signed rank test was used in the case of same-cell experiments. Two-tailed log rank analyses were used to analyze statistical significance of Kaplan–Meier survival curves. A level of P < 0.05 was used to designate significant differences. Based on the variance of xenograft growth in control mice, we used at least three mice per genotype to give 80% power to detect an effect size of 20% with a significance level of 0.05. For all mouse experiments, the number of independent mice used is listed in figure legend. Statistical analyses of retrospective patient data are described above.

## Data availability

All data are available in the manuscript or from the corresponding author upon reasonable request. Source data will be uploaded with the final version of the manuscript.

## Code availability

Sources for all code used have been provided, no custom code was created for this manuscript.

## Acknowledgements

This work was supported by grants from Cancer Research UK (to M.M.), ChadTough Defeat DIPG (to M.M. and T.B.), the National Institute of Neurological Disorders and Stroke (R01NS092597 to M.M.), NIH Director’s Pioneer Award (DP1NS111132 to M.M.), National Cancer Institute (P50CA165962, R01CA258384, U19CA264504 to M.M.), Robert J. Kleberg, Jr. and Helen C. Kleberg Foundation (to M.M.), McKenna Claire Foundation (to M.M.), Kyle O’Connell Foundation (to M.M.), Virginia and D.K. Ludwig Fund for Cancer Research (to M.M.), Waxman Family Research Fund (to M.M.), Will Irwin Research Fund (to M.M.). The authors thank Shawn Hervey-Jumper for the gift of IDH WT adult GBM SF0232 and SF0238 cells.

## Author contributions

M.M. and T.B. designed the experiments and wrote the manuscript. T.B., B.Y., K.S. V.M., S.M.J., K.R.T, M.B.K., H.X., M.S., M.A.Q., and P.J.W conducted experiments and performed data analyses. P.G.F., S.P., C.J.C., A.M., C.K., D.L. maintained patient databases at Stanford and University of Michigan; A.M., E.C., A.F., S.L., abstracted data from the databases; D.T. and C.K. reviewed source data for all pediatric high-grade glioma database entries to ensure veracity and completeness.

All authors contributed to manuscript editing. M.M. conceived the project and supervised all aspects of the work.

## Author declarations

M.M. holds equity in MapLight Therapeutics and Syncopation Life Sciences.

---

**Extended Data Figure 1.**
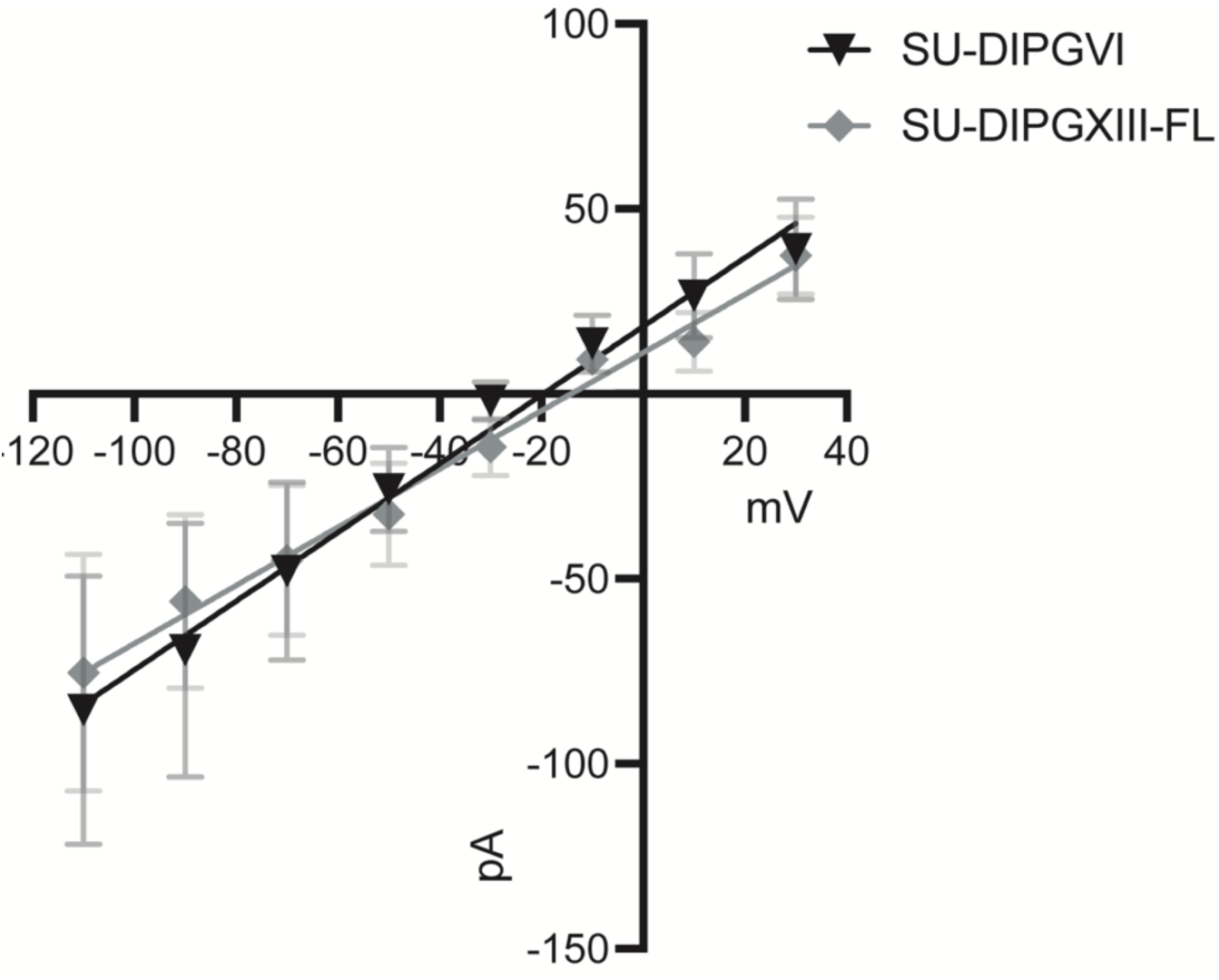
Current-voltage relationship of GABA current in two patient-derived DMG xenograft models recorded with perforated patch. Reversal potential of GABA was -19.61 mV in SU-DIPGVI cells (n = 6 cells from 5 mice), and -14.14 mV in SU-DIPGXIII-FL cells (n=5 cells from 3 mice). All data are mean ± s.e.m.

**Extended Data Figure 2.**
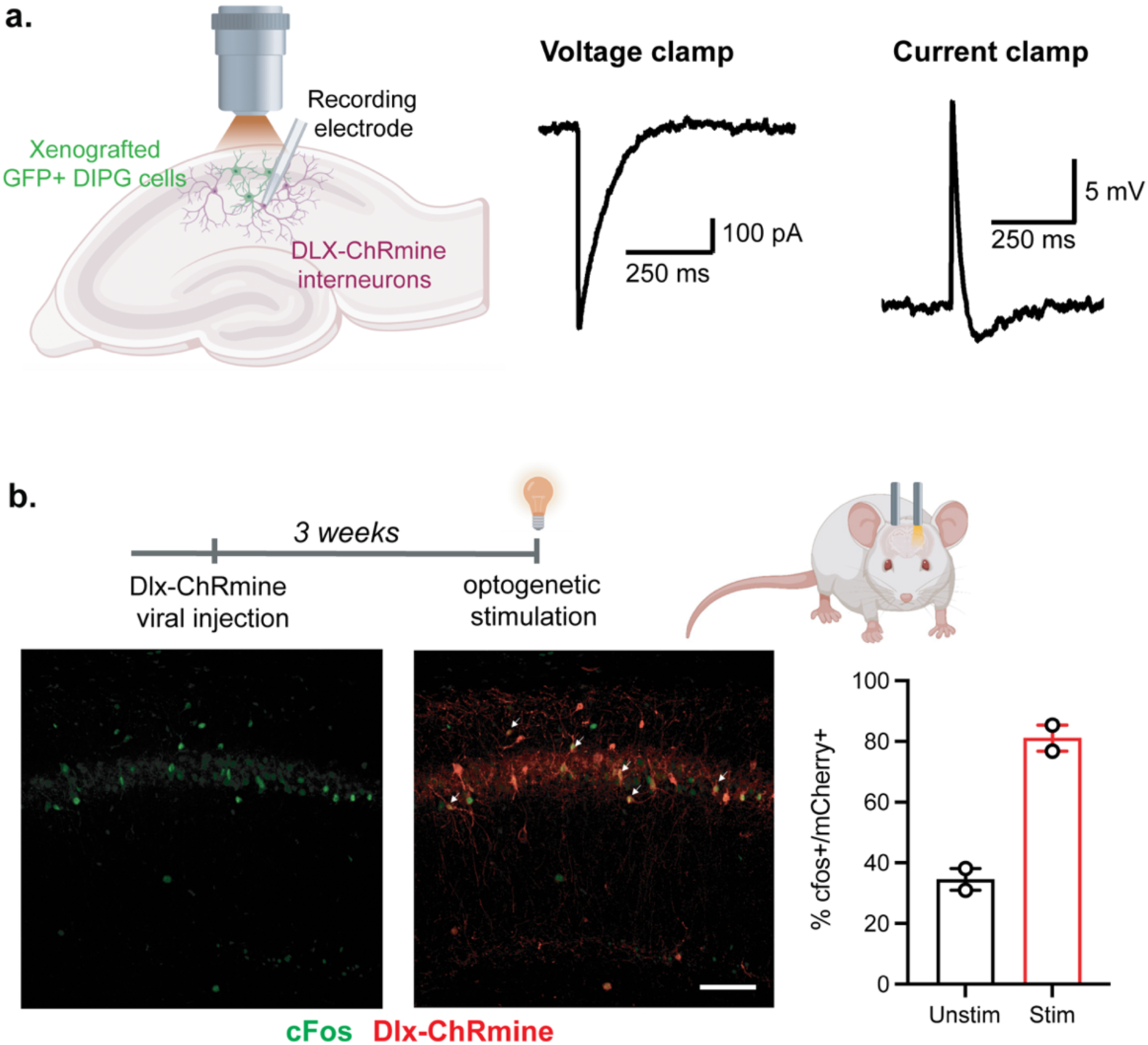
Optogenetic stimulation of GABAergic interneurons expressing DLX-ChRmine. **a.** Inward current and corresponding depolarization of GABAergic interneurons expressing DLX-ChRmine in response to optogenetic stimulation were recorded in using patch clamp electrophysiology. **b.** Optogenetic stimulation of interneurons expressing DLX-ChRmine (red) lead to neuronal activity, indicated by cfos expression (green). Arrows indicate co-labeled cells. Scale bar, 100 µm. Above, experimental timeline. Right, quantification of cfos expression in interneurons in the stimulated hemisphere (stim) is greater than in the unstimulated hemisphere (unstim, n = 2 mice). All data are mean ± s.e.m.

**Extended Data Figure 3.**
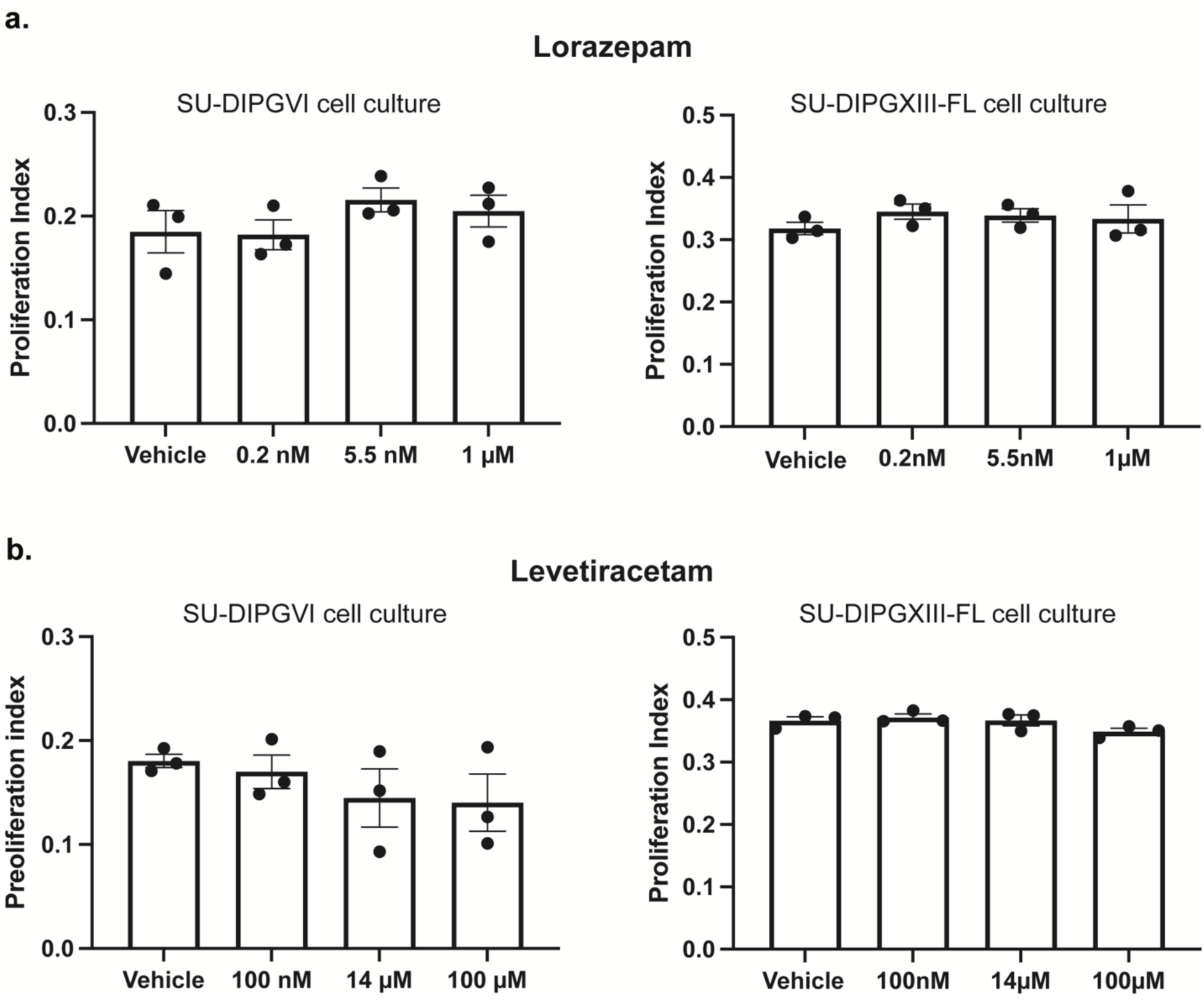
Lorazepam and levetiracetam have no effect on proliferation of patient-derived DMG cells in monoculture. **a.** Lorazepam treatment in patient-derived DMG cultures SU-DIPGVI and SU-DIPGXIII-FL had no effect on proliferation (n = 3 wells per group). **b.** Levetiracetam treatment in patient-derived DMG cultures SU-DIPGVI and SU-DIPGXIII-FL had no effect on proliferation (n = 3 wells per group). All data are mean ± s.e.m. Two-tailed Student’s t-test.

**Extended Data Figure 4.**
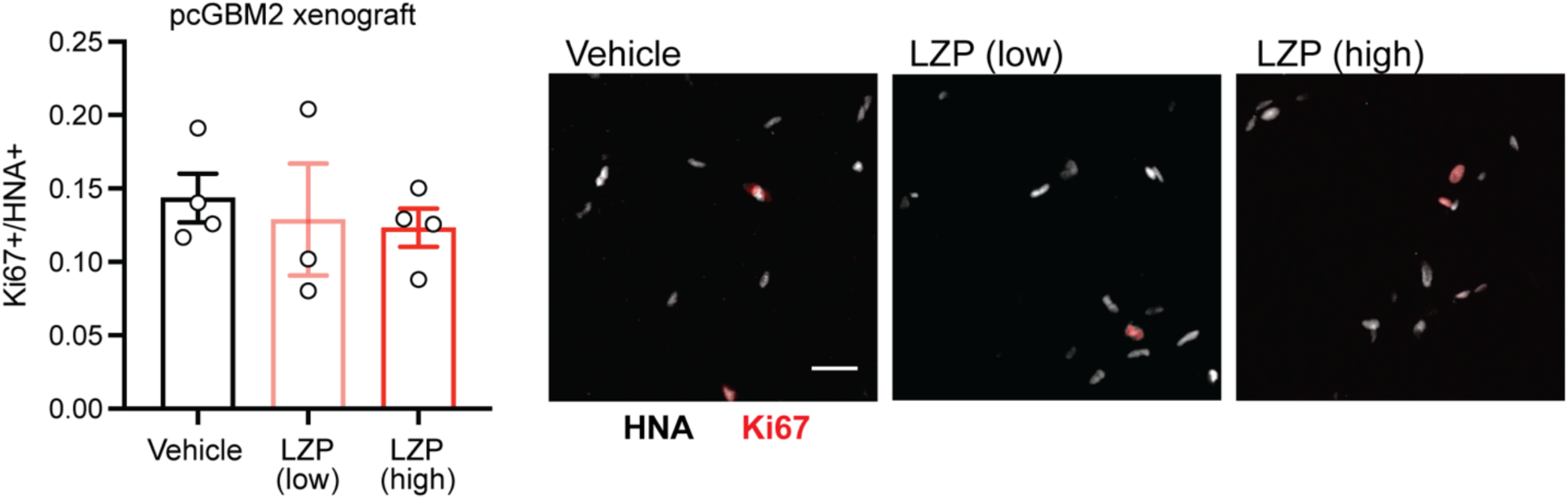
Lorazepam has no effect on H3/IDH WT pediatric GBM. LZP treatment in mice with patient-derived pcGBM2 xenografts has no effect on cell proliferation (vehicle, n = 4 mice; low dose, n = 3 mice; high dose, n = 4 mice). Representative confocal images of xenografted pcGBM2 cells expressing Ki67 (red) and HNA (white; right). Scale bar, 25 µm. All data are mean ± s.e.m. One-way ANOVA.

**Extended Data Figure 5.**
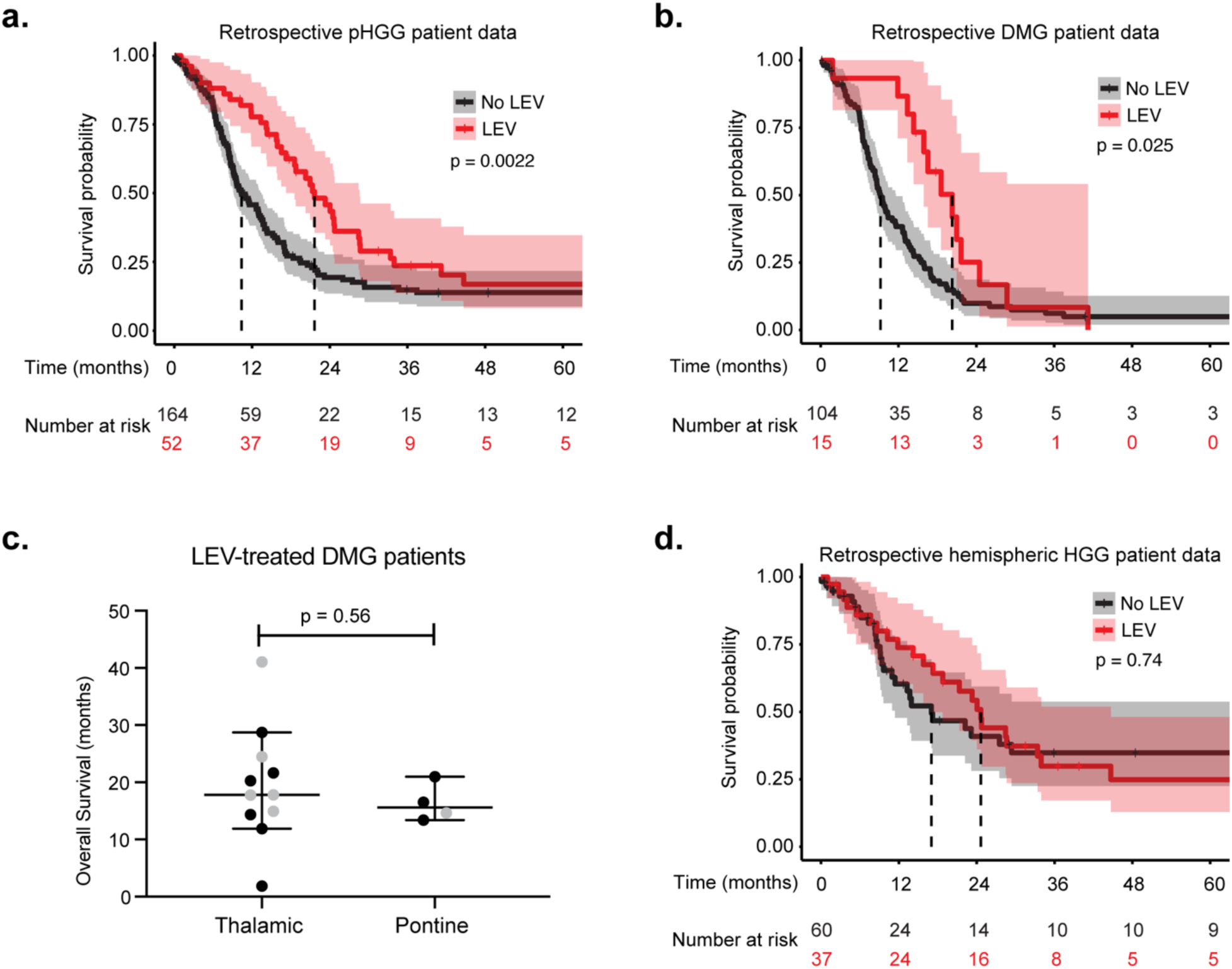
Effect of levetiracetam on overall survival in retrospective, real-world patient data. **a.** All pediatric high-grade gliomas: Kaplan–Meier overall survival (OS) curves of retrospective data from Stanford University (1990-2020) and University of Michigan (2012-2021) patient databases (n=216), showing pediatric high-grade glioma (pHGG) patients treated with LEV (LEV: median OS = 21.7 months, n = 52; no LEV: median OS= 10.4 months, n = 164). **b.** Diffuse midline gliomas: Kaplan–Meier overall survival (OS) curves of retrospective data from DMG patients treated with LEV (LEV: median OS = 20.3 months, n = 15; no LEV: median OS= 9.2 months, n = 105). **c.** No difference in overall survival of LEV-treated patients with thalamic compared to pontine DMG (thalamic: n = 11; pontine: n = 4). Grey points are data censored at time of last follow-up. Data are median ± 95% CI. n.s. (P>0.05), two-tailed Student’s t-test. **d.** Hemispheric high-grade gliomas: Kaplan–Meier overall survival (OS) curves of retrospective data from hemispheric high-grade glioma (HGG) patients treated with LEV (LEV: OS=24.6 months, n = 37; no LEV: OS = 17.0 months, n = 60).

**Extended Data Figure 6.**
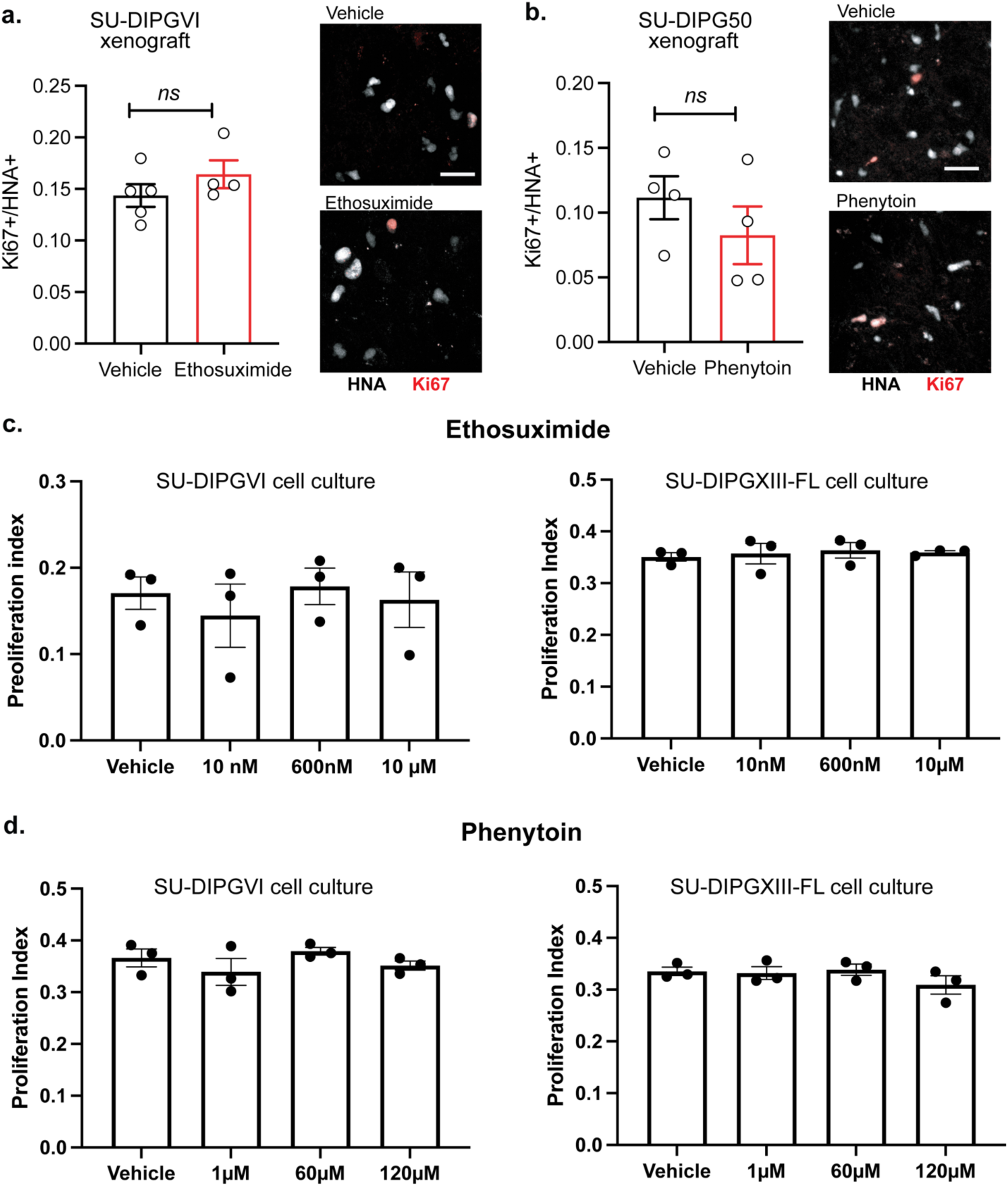
Antiepileptic drugs ethosuximide and phenytoin do not affect DMG proliferation. **a.** Ethosuximide treatment in mice with patient-derived SU-DIPGVI xenografts has no effect on cell proliferation (vehicle, n = 5 mice; ethosuximide, n = 4 mice). Representative confocal images of xenografted SU-DIPGVI cells expressing Ki67 (red) and HNA (white; right). Scale bar, 25 µm. **b.** Phenytoin treatment in mice with patient-derived SU-DIPG50 xenografts has no effect on cell proliferation (vehicle, n = 4 mice; phenytoin, n = 4 mice). Representative confocal images of xenografted SU-DIPG50 cells expressing Ki67 (red) and HNA (white; right). Scale bar, 25 µm. **c.** Ethosuximide treatment in patient-derived DMG cultures SU-DIPGVI and SU-DIPGXIII-FL had no effect on proliferation (n = 3 wells per group). **d.** Phenytoin treatment in patient-derived DMG cultures SU-DIPGVI and SU-DIPGXIII-FL had no effect on proliferation (n = 3 wells per group). All data are mean ± s.e.m. Two-tailed Student’s t-test.

**Extended Data Table 1.**
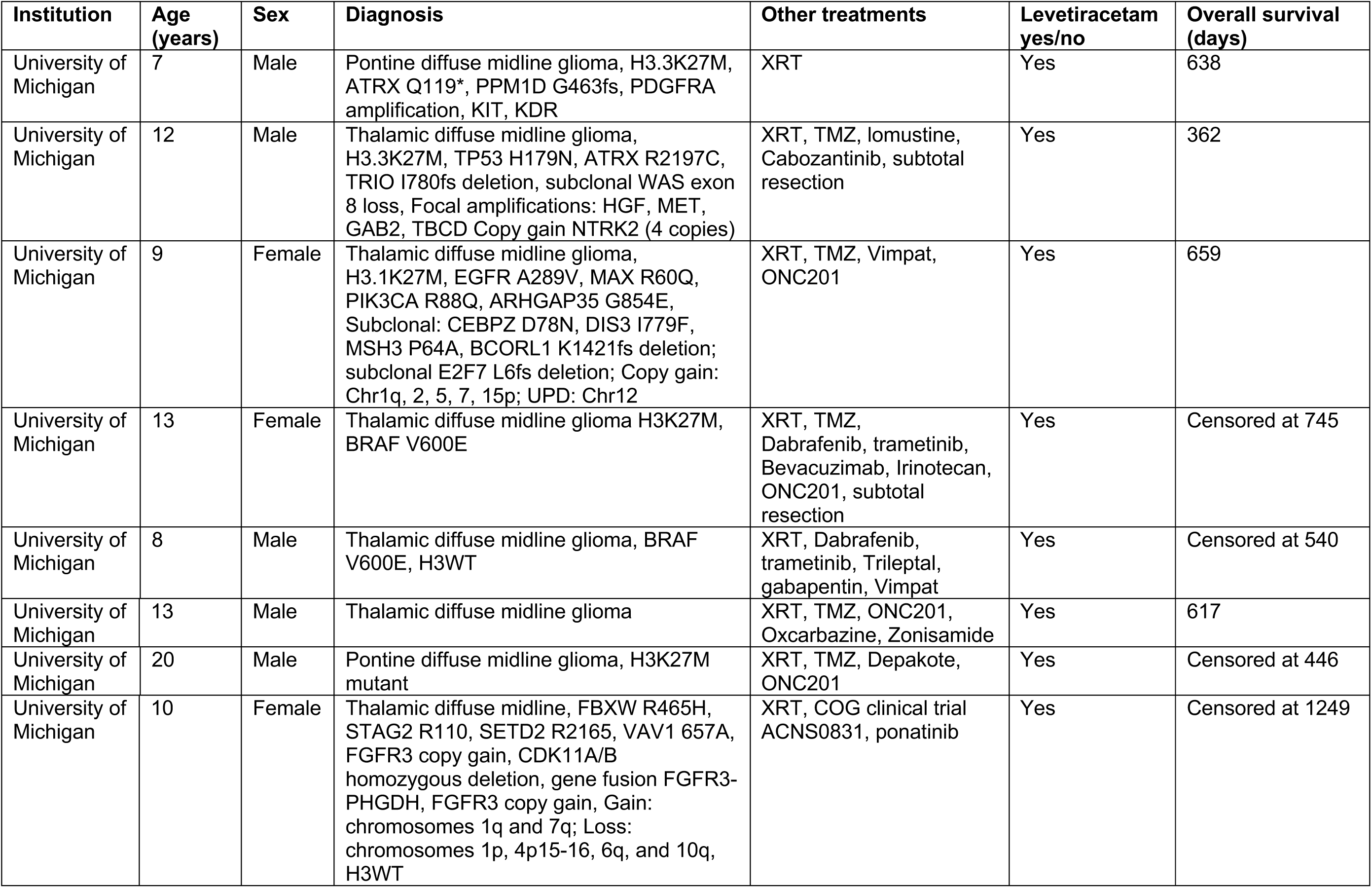
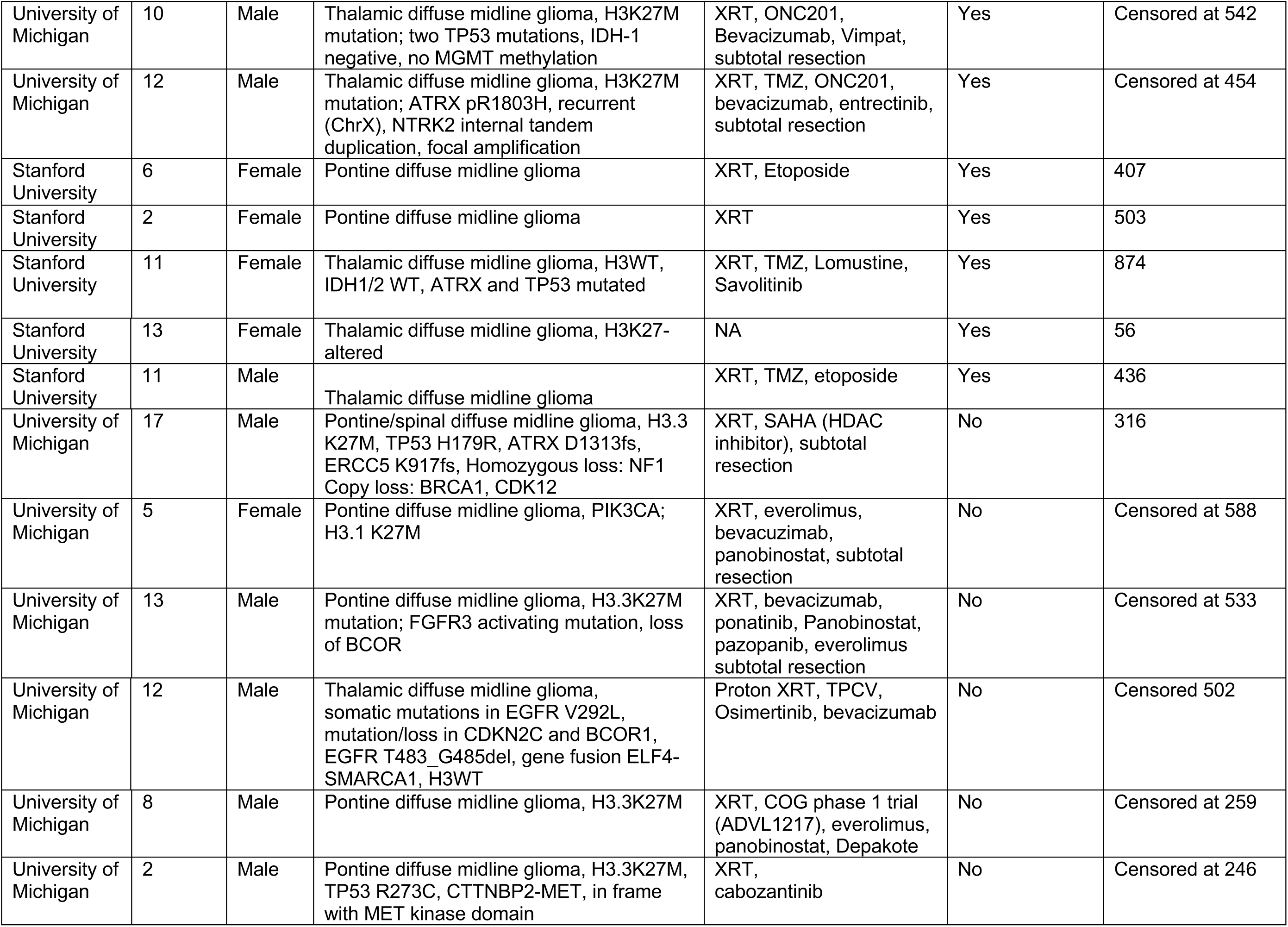
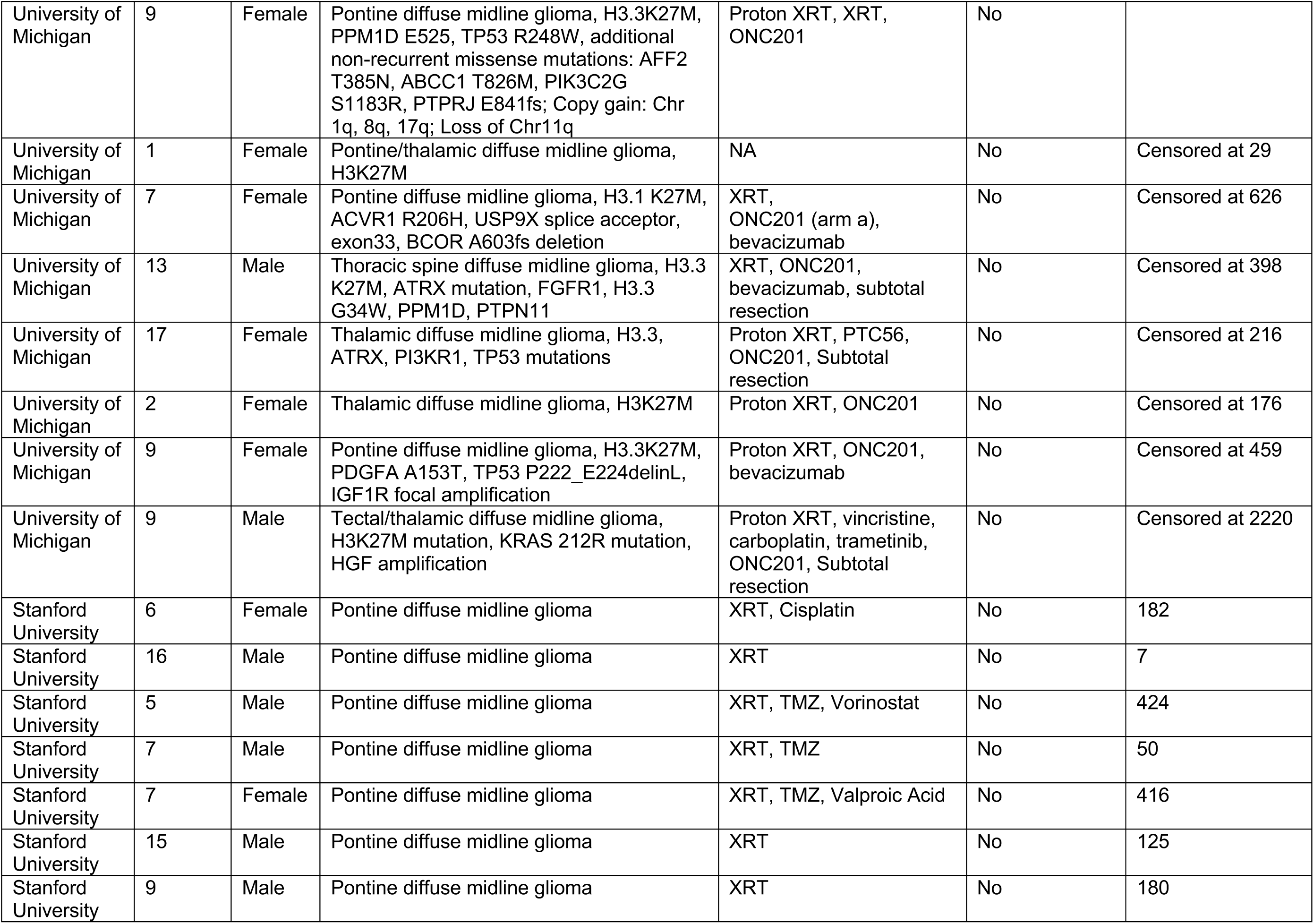
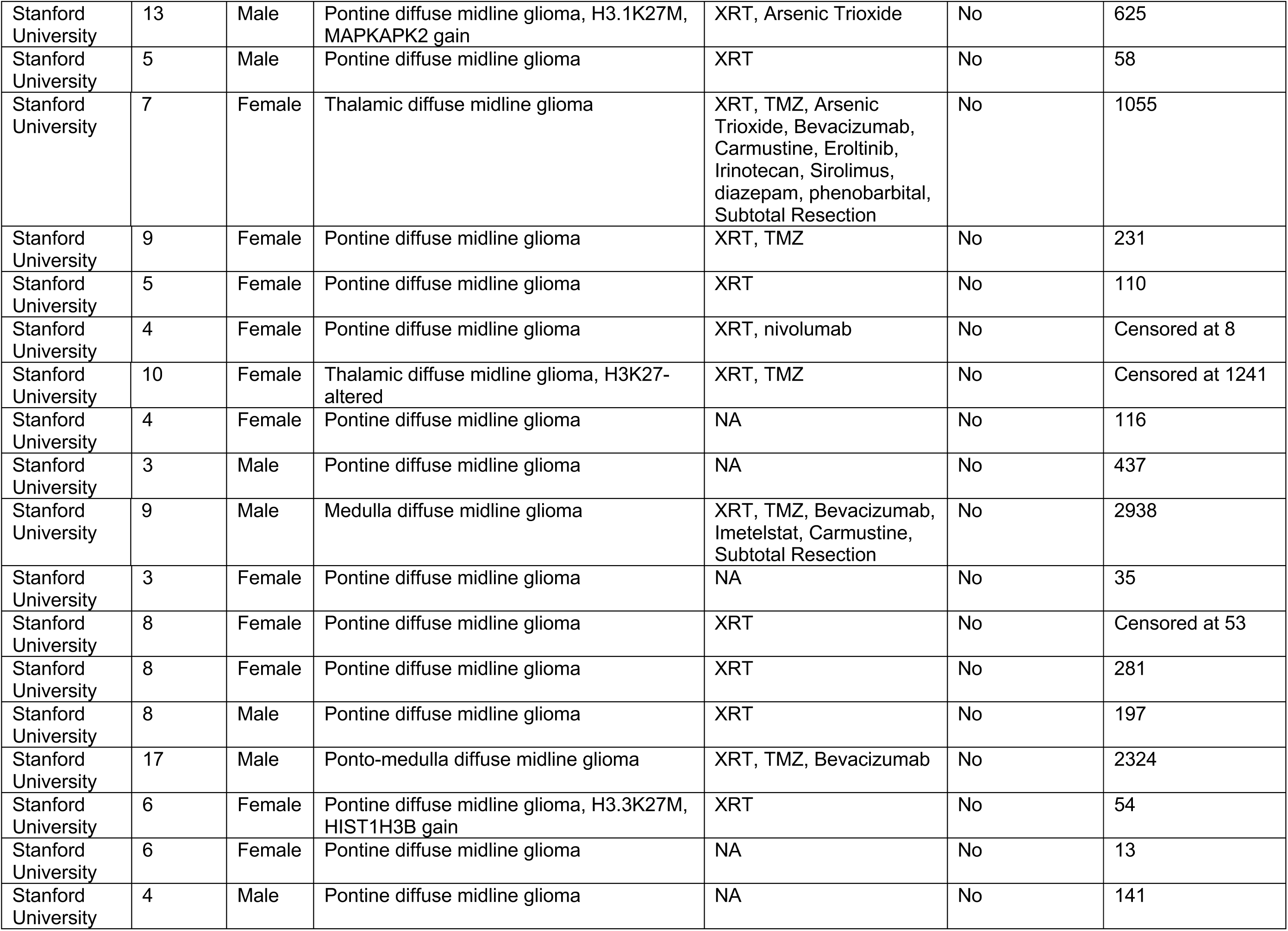
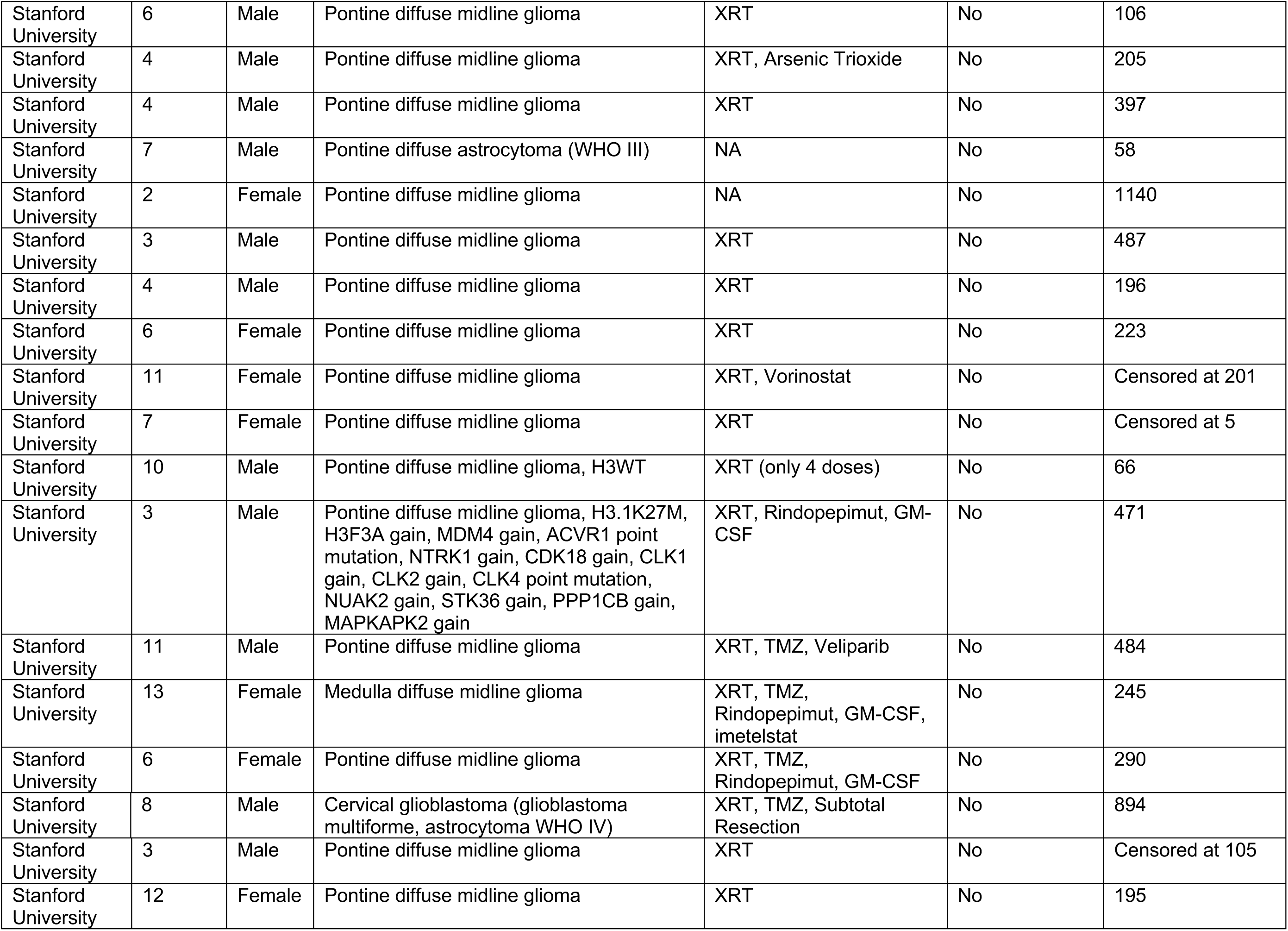
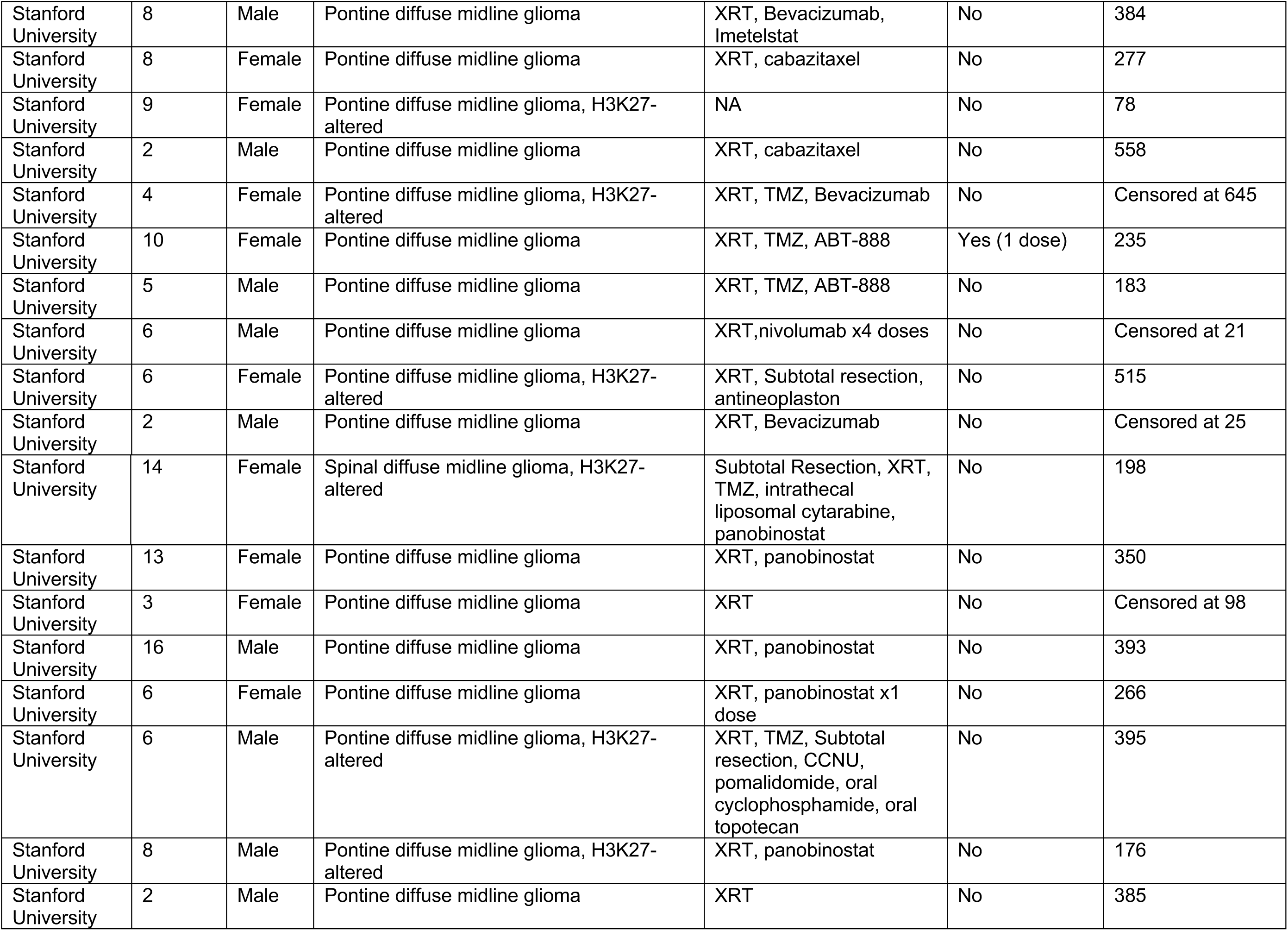
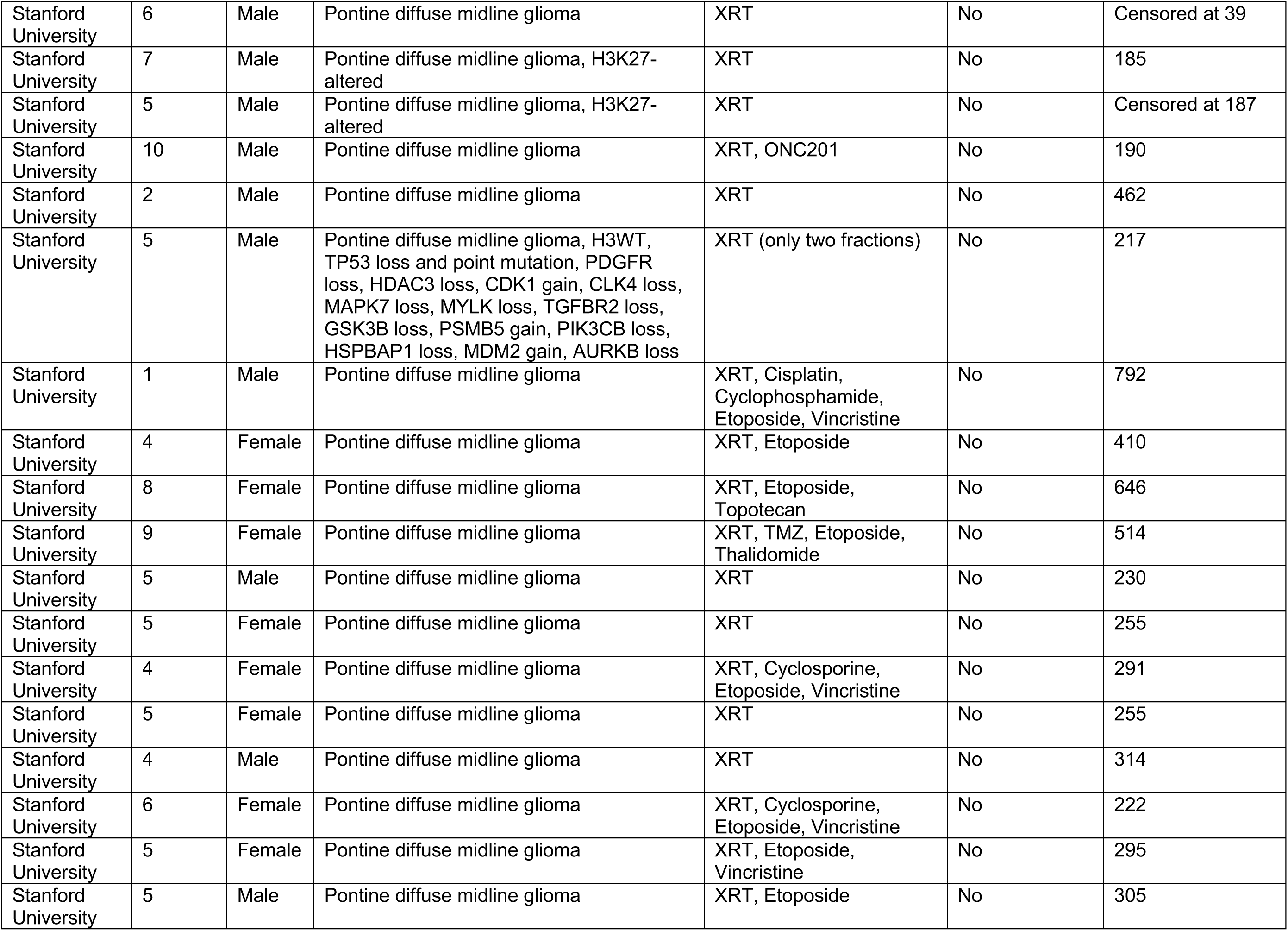
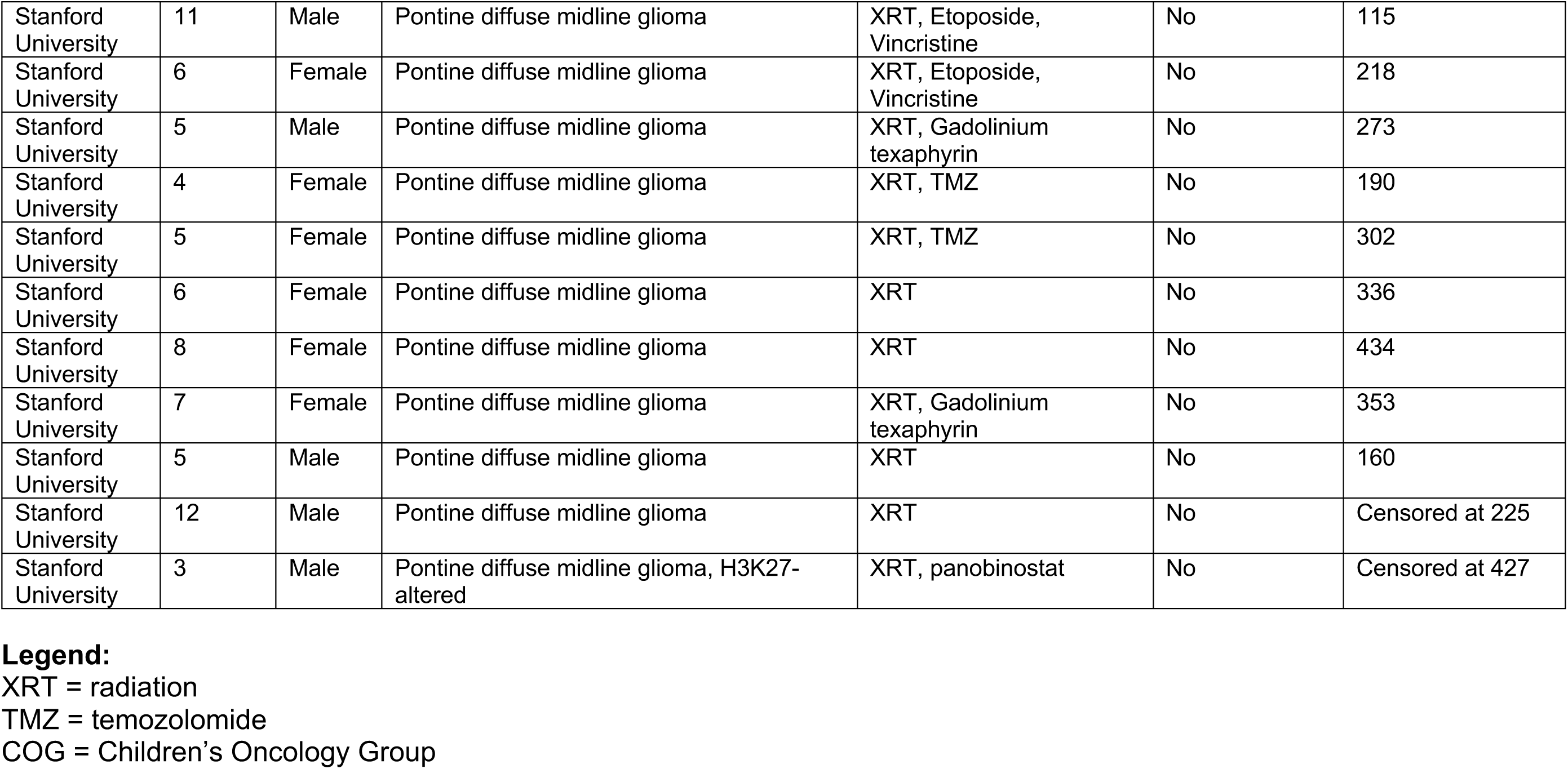
Diffuse Midline Glioma Patients

**Extended Data Table 2.**
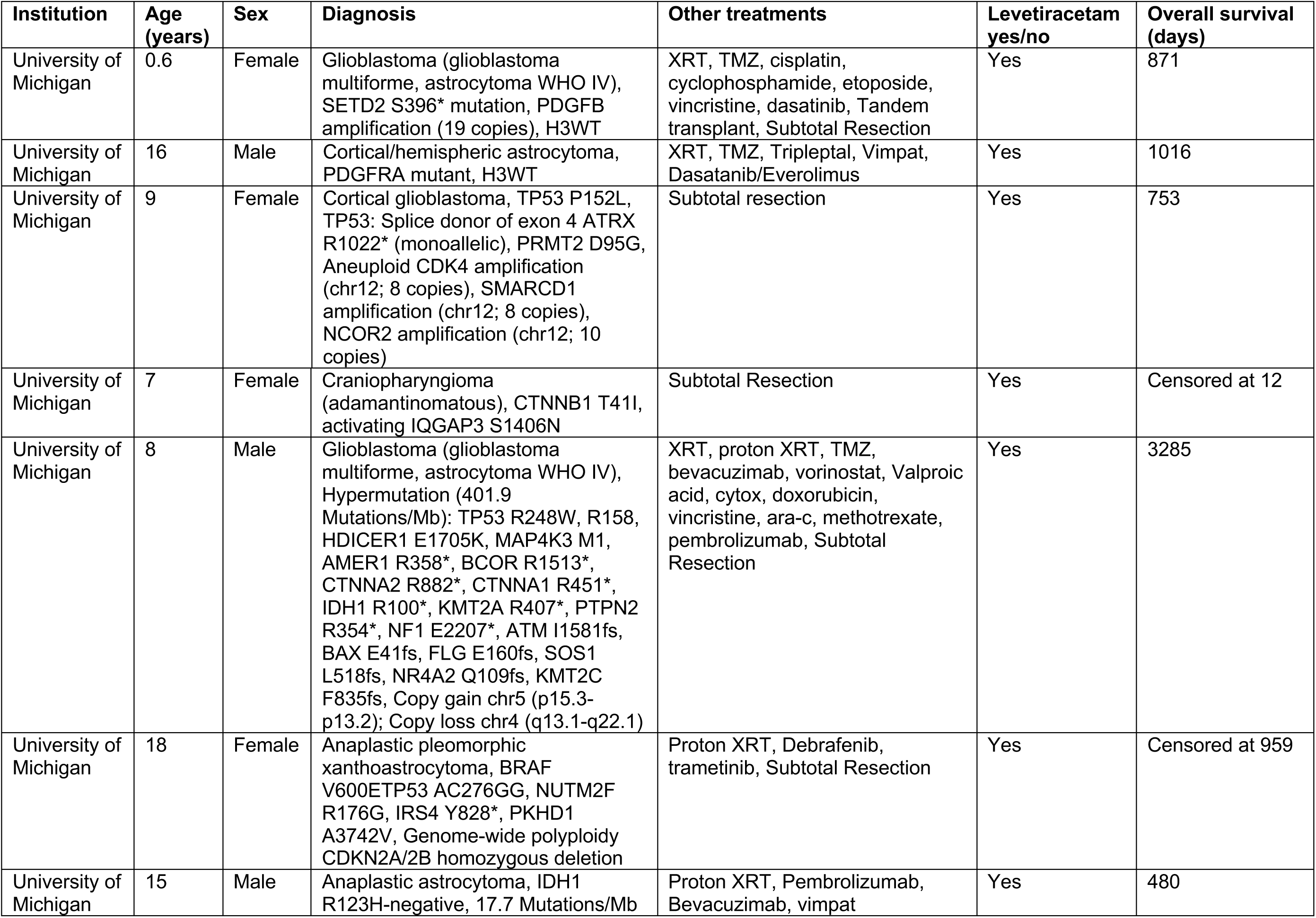
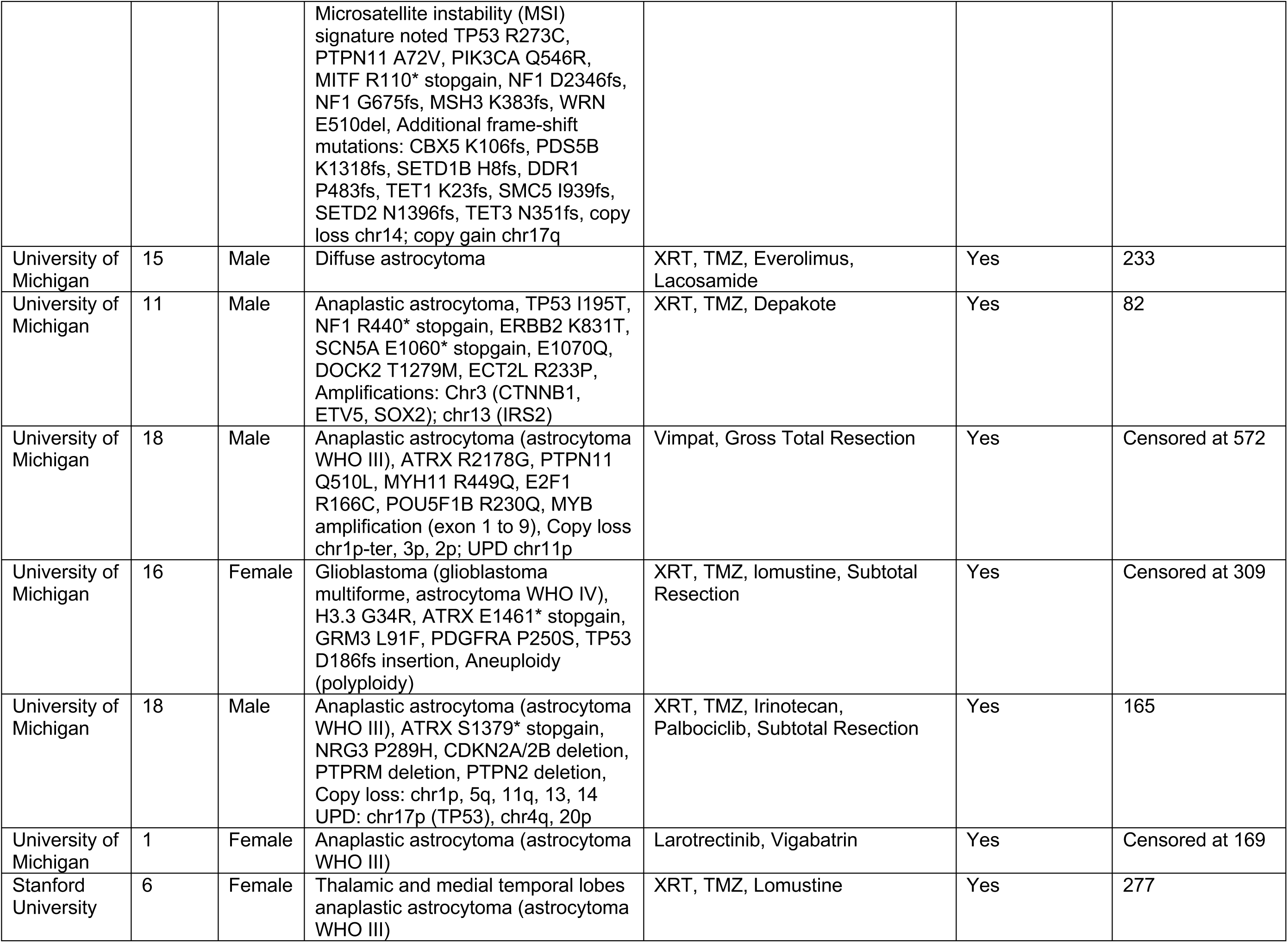
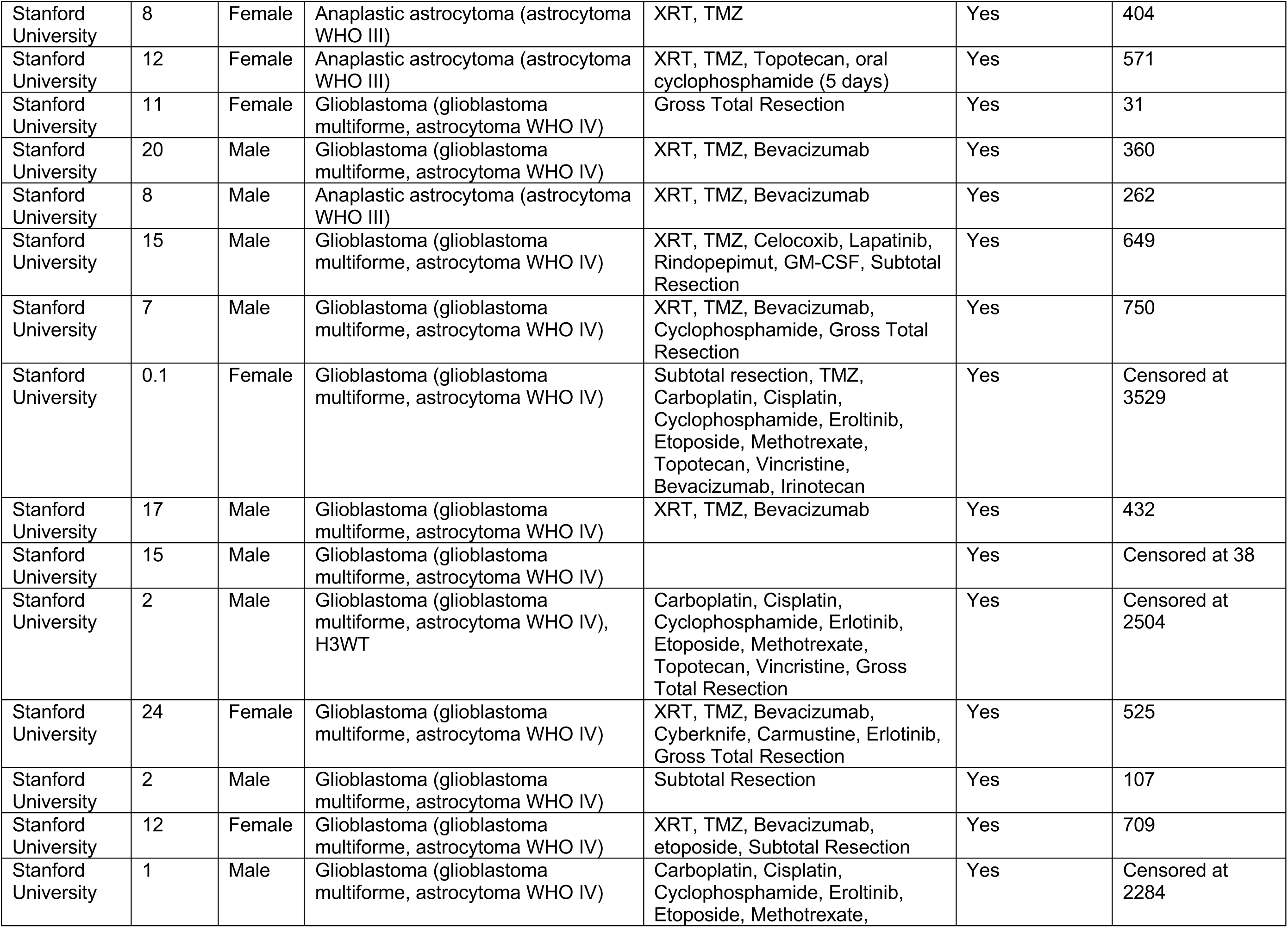
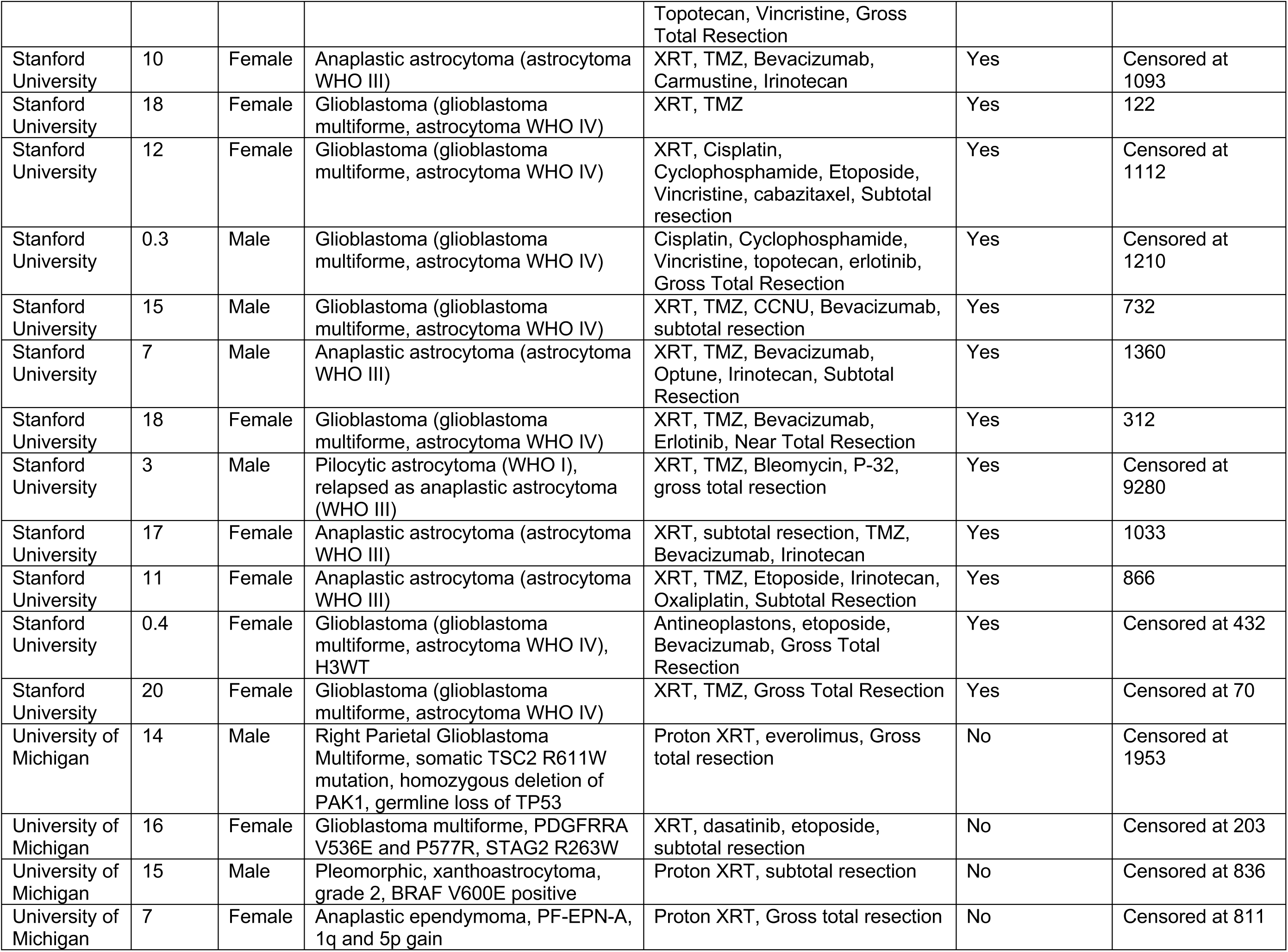
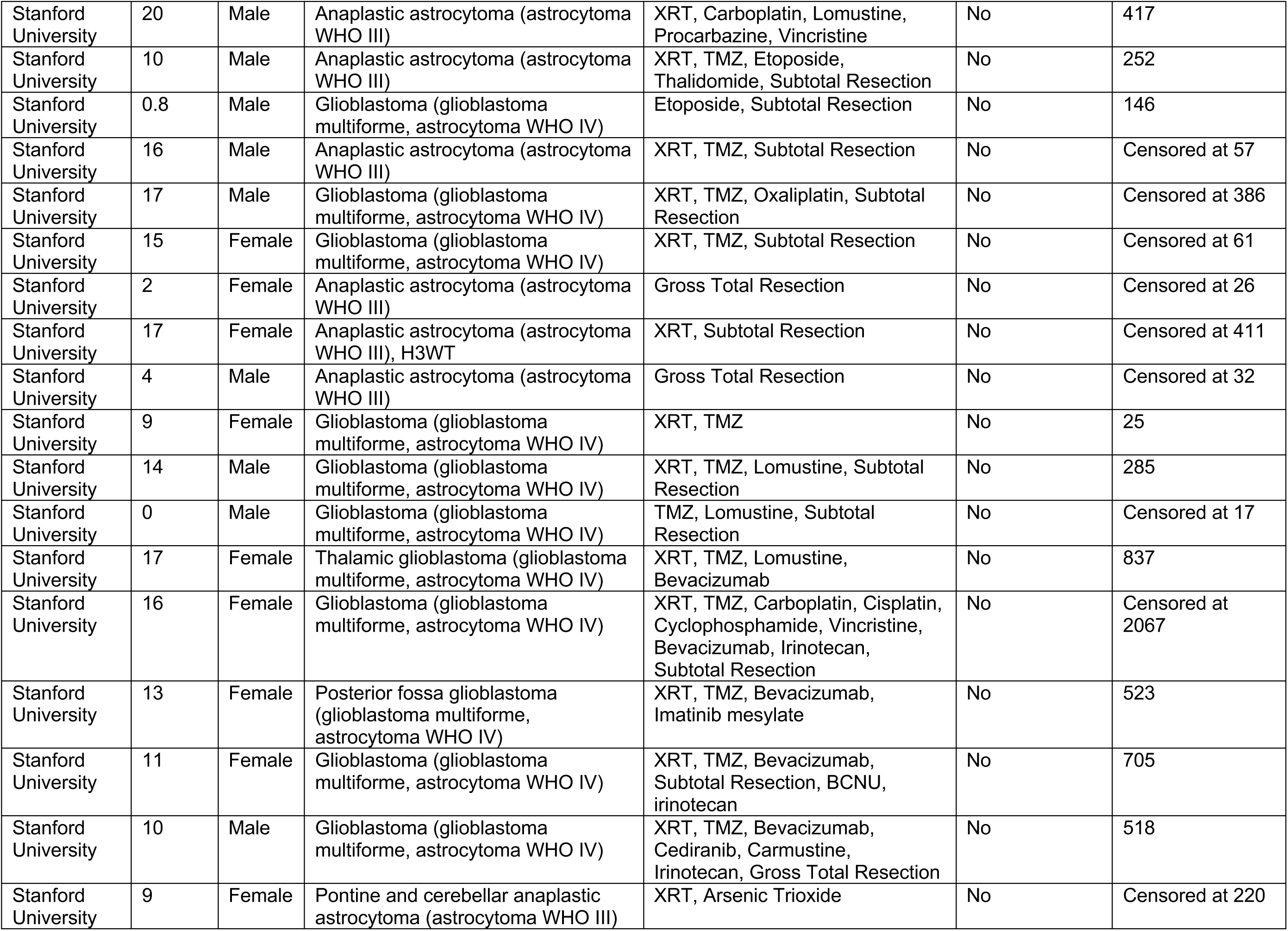
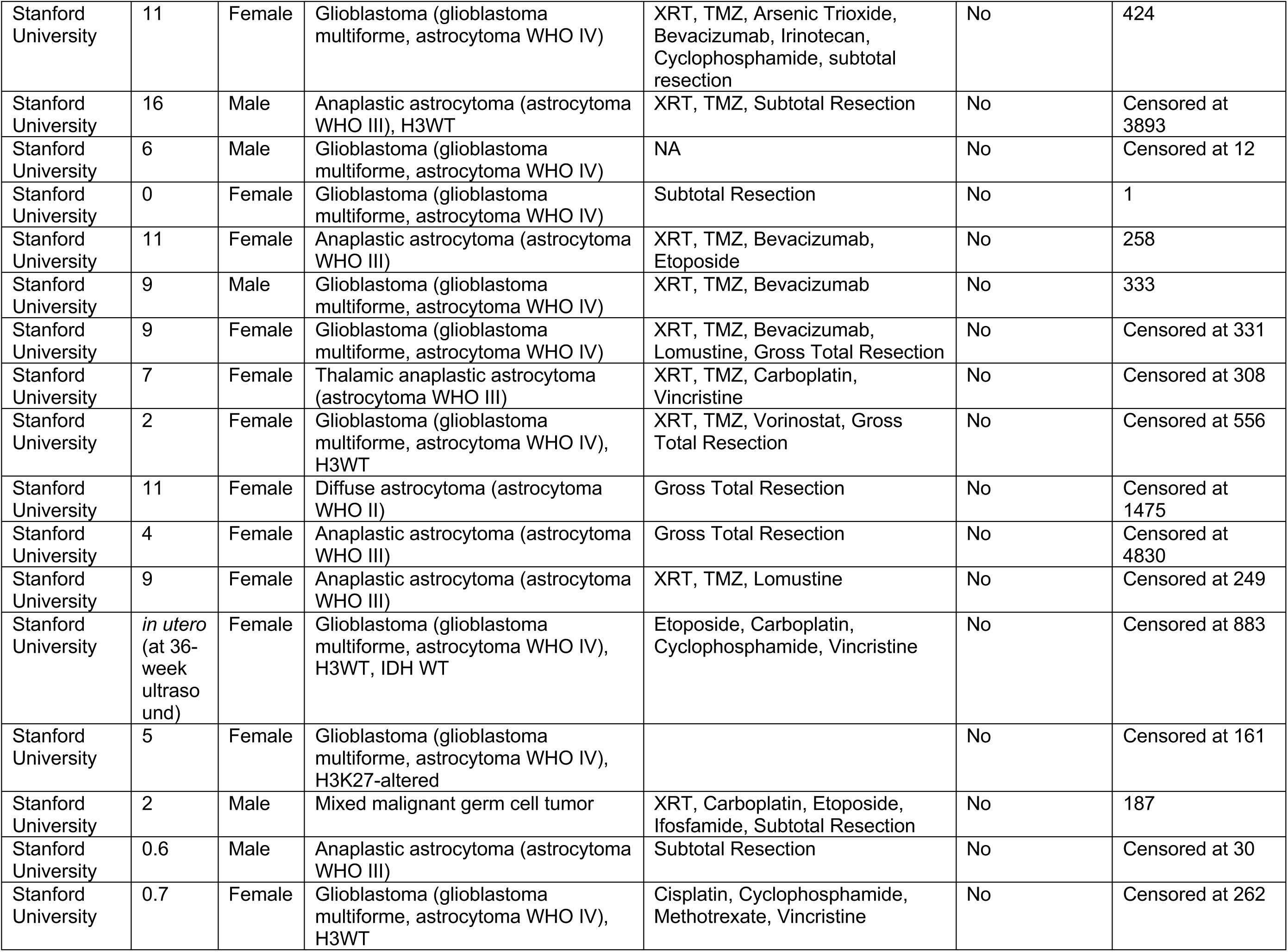
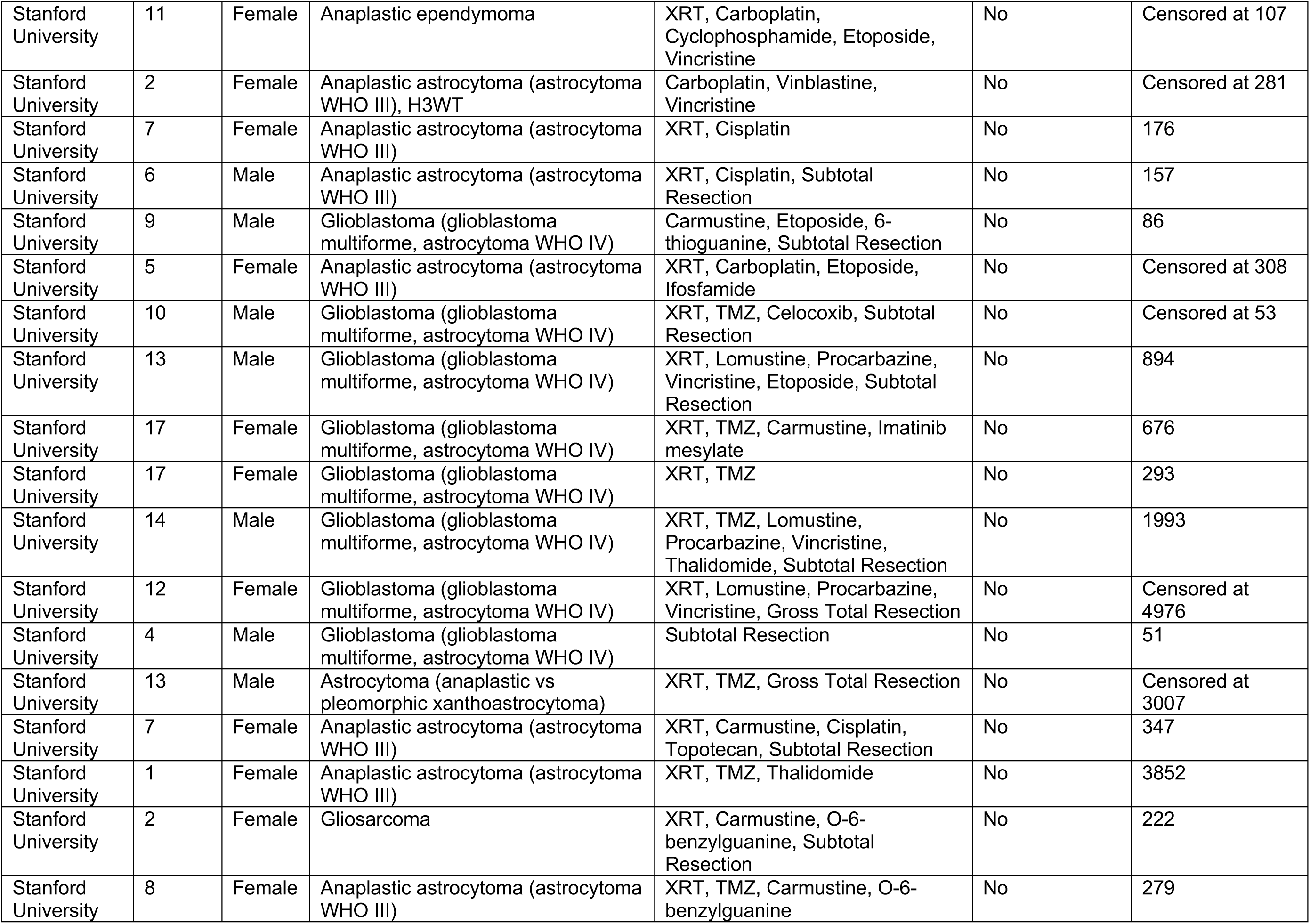
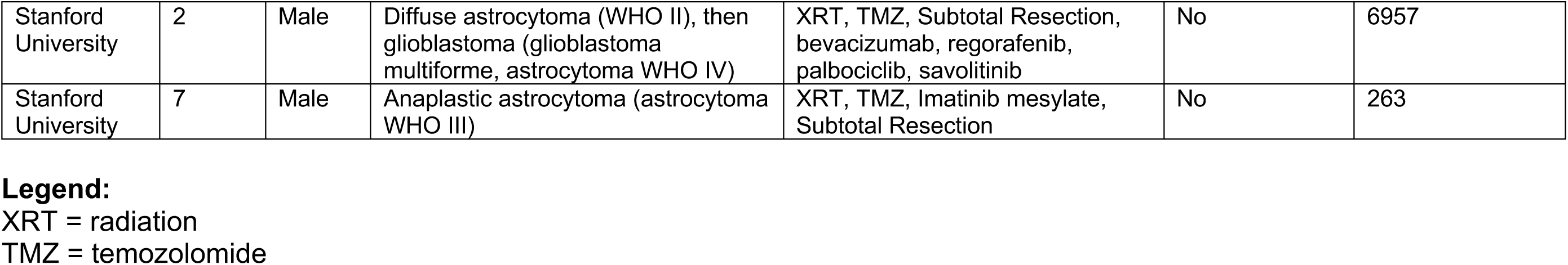
Hemispheric High Grade Glioma Patients

